# The action mechanism of actinoporins revealed through the structure of pore-forming intermediates

**DOI:** 10.1101/2024.06.27.601005

**Authors:** Rocío Arranz, César Santiago, Simonas Masiulis, Esperanza Rivera-de-Torre, Juan Palacios-Ortega, Diego Carlero, Diego Heras-Márquez, José G. Gavilanes, Ernesto Arias-Palomo, Álvaro Martínez-del-Pozo, Sara García-Linares, Jaime Martín-Benito

**Affiliations:** Departamento de Estructura de Macromoléculas, Centro Nacional de Biotechnología (CNB-CSIC), 28049, Madrid, Spain; Materials and Structural Analysis Division, Thermo Fisher Scientific, Achtseweg Noord 5, 5651, Eindhoven, The Netherlands; Departamento de Bioquímica y Biología Molecular, Universidad Complutense, Madrid, Spain; Centro de Investigaciones Biológicas Margarita Salas, CSIC, 28040 Madrid, Spain

**Author notes:** Corresponding authors: SGL and JMB. Department of Biotechnology and Biomedicine, Technical University of Denmark, Kongens Lyngby, Denmark (Esperanza Rivera-de-Torre). Department of Chemistry, Faculty of Natural, Mathematical & Engineering Sciences, King’s College London, London, United Kingdom (Juan Palacios-Ortega). These authors contributed equally to this work (Rocío Arranz, César Santiago, Simonas Masiulis).

## Abstract

Pore-forming proteins exemplify the transformative potential of biological molecules. Initially produced in a monomeric, water-soluble form, they spontaneously assemble into multimeric integral membrane proteins in the presence of suitable target lipids. Their functions include roles in apoptosis, cell signaling, immunity, as well as attack and defense systems between different organisms. This latter group encompasses actinoporins, a family of pore-forming toxins from sea anemones that kill target cells by perforating their plasma membrane. Here, we have determined the structures of two such toxins, fragaceatoxin C and sticholysin II, in a membrane environment using cryogenic electron microscopy. The structures reveal how dozens of lipid molecules interact in an orderly manner, forming an intrinsic part of the pore. We have also isolated different pore-forming intermediates, where only a fraction of the constituent monomers is incorporated, exhibiting non-closed, arc-shaped structures. Based on these structures we propose a mechanism of action where the sequential assembly of toxin monomers onto the membrane, accompanied by conformational changes, triggers pore formation and membrane perforation. Our results contribute to a better understanding of the transforming capacity of these pore-forming proteins, which are becoming increasingly important for their diverse biotechnological applications.

## Introduction

Pore-forming proteins are a rather unique group of metamorphic proteins synthesized as water-soluble monomers which, in contact with the appropriate lipid bilayer, oligomerize and convert into integral membrane proteins to build a pore [1]. The functions of these proteins vary widely, encompassing, among others, programmed cell death, signaling, immunity, or toxins that attack other cells or organisms. Pore-forming toxins (PFTs) constitute the major class within the pore-forming proteins, and have been found in viruses and bacteria, and higher organisms [2, 3]. Depending on the secondary structure constituting the transmembrane part of the pore, these toxins are classified either as α- or β-PFTs [4–6]. Among the α-PFTs are the actinoporins, a family of toxins of ∼20kDa molecular weight synthesized by sea anemones. The study of these proteins is of increasing interest not only because they constitute an optimal system to investigate the water-membrane transition of proteins, but also because these proteins are indeed proving to be an ideal platform for building new and sustainable biotechnological tools with improved functionalities, such as DNA sequencing or bioremediation [7–9].

All known actinoporins adopt a well-conserved water-soluble three-dimensional fold (Extended Data Fig. 1). This monomeric structure consists of a β-sandwich, made of 10-12 β-strands, flanked by two α-helices [10–13]. One of the α-helices is located at the N-terminal end (α-helix 1) and seems to be the only structural element undergoing a significant transformation when the protein binds to the membrane [14–20]. The incorporation of actinoporins into a membrane, and the subsequent formation of a pore, depends largely on the lipid composition and its derived physicochemical properties [21–30]. Sphingomyelin (SM) is specifically required [24, 31–33], but other conditions have a strong influence on their pore-forming ability. Some examples of these dependences include the coexistence of various phases or domains, lateral packing, fluidity, membrane thickness, the strength of the interfacial hydrogen bonding network, or the presence of sterols [18, 24, 29, 34–39]. In fact, it is well-known that cholesterol (Chol) greatly influences the pore-forming behavior [26, 29, 37, 38, 40–44], although the molecular details still remain unknown. Once bound to the membrane, the most generally accepted mechanism of pore formation suggests that α-helix 1 extends to the first ∼30 residues and then it is inserted into the membrane to form the pore walls [13, 15–17, 34, 41, 45, 46]. This conformational change has to occur in coordination with the oligomerization needed to assemble a stable pore, but the details of this mechanism are far from being understood.

Here, using cryogenic electron microscopy (cryoEM), we have determined the structures of two actinoporins, sticholysin II (StnII) from *Stichodactyla helianthus* [47] and fragaceatoxin C (FraC) from *Actinia fragacea* [48], within the membrane environment using large unilamelar vesicles (LUVs) and lipid nanodiscs [49], both systems containing 1,2-dioleoyl-sn-glycero-3-phosphocholine (DOPC), porcine brain SM, and Chol. Our findings reveal three key aspects of the pore assembly process. Firstly, lipids play a pivotal role in the final assembly, with SM and Chol forming two well-structured rings containing up to eighty-eight molecules surrounding the pore. Secondly, while different oligomerization stoichiometries are possible, the octameric arrangement is the most abundant and thermodynamically stable. And thirdly, it is possible to isolate pore-assembling intermediates in lipid nanodiscs. Based on these results, we propose a molecular mechanism that outlines the specific sequential events leading to the pore formation.

## Results

### Structure of the StnII and FraC oligomeric pores on lipid membranes

To elucidate the structure of the FraC and StnII in their natural membrane environment, we used both liposomes and lipid nanodiscs. The pores were obtained by mixing the water-soluble protein with either LUVs or nanodiscs of the composition DOPC:SM:Chol (80:20:10), as described in the methods section.

Liposome samples revealed that the formed pores tend to cluster in patches when the membrane is not fully covered, leaving certain regions of the vesicle bare (Extended Data Fig. 2A). The pore clusters increase the membrane curvature, producing dome-shaped deformations that protrude from the liposome. At higher toxin concentrations, liposomes are fully covered by pores and exhibit more uniform spherical shapes and reduced diameters, indicating that liposome cleavage may have occurred. On the other hand, the nanodisc preparations are prone to exhibit a preferred orientation on the grid, mostly showing views along the pore longitudinal axis (“top views”) and, to a lesser extent, views perpendicular to the longitudinal axis (“side views”) (Extended Data Fig. 2B). These behaviors are common to both FraC and StnII proteins.

The structure of FraC in liposomes, resolved at 2.1 Å resolution, reveals an octameric pore with a funnel-shaped arrangement, where the widest part corresponds to the extracellular region when the protein is functioning in its natural environment (Figure 1, Extended Data Fig. 3). The structure closely resembles the one described by Tanaka et al., which was obtained using detergents and X-ray crystallography [13] (Cα RMSD = 0.388 Å). However, our data provide a more detailed representation of organization of the lipids forming part of the pore because no detergents were used, allowing us to identify up to eighty-eight molecules surrounding the pore. The lipids are located in the middle and upper parts of the transmembrane helix that forms the pore walls and are distributed as seven SM, and four Chol molecules per protein monomer. Three of these seven SMs coincide with the positions defined as L1 to L3 in the X-ray structure (4TSY, [13]), including the lipid molecule present in the fenestration. However, we were able to determine the acyl chains of the lipids due to the higher resolution of the map (Figure 2). Furthermore, there are two extra acyl chains in the lower part of the transmembrane region (Figure 1C).

**Figure 1.**
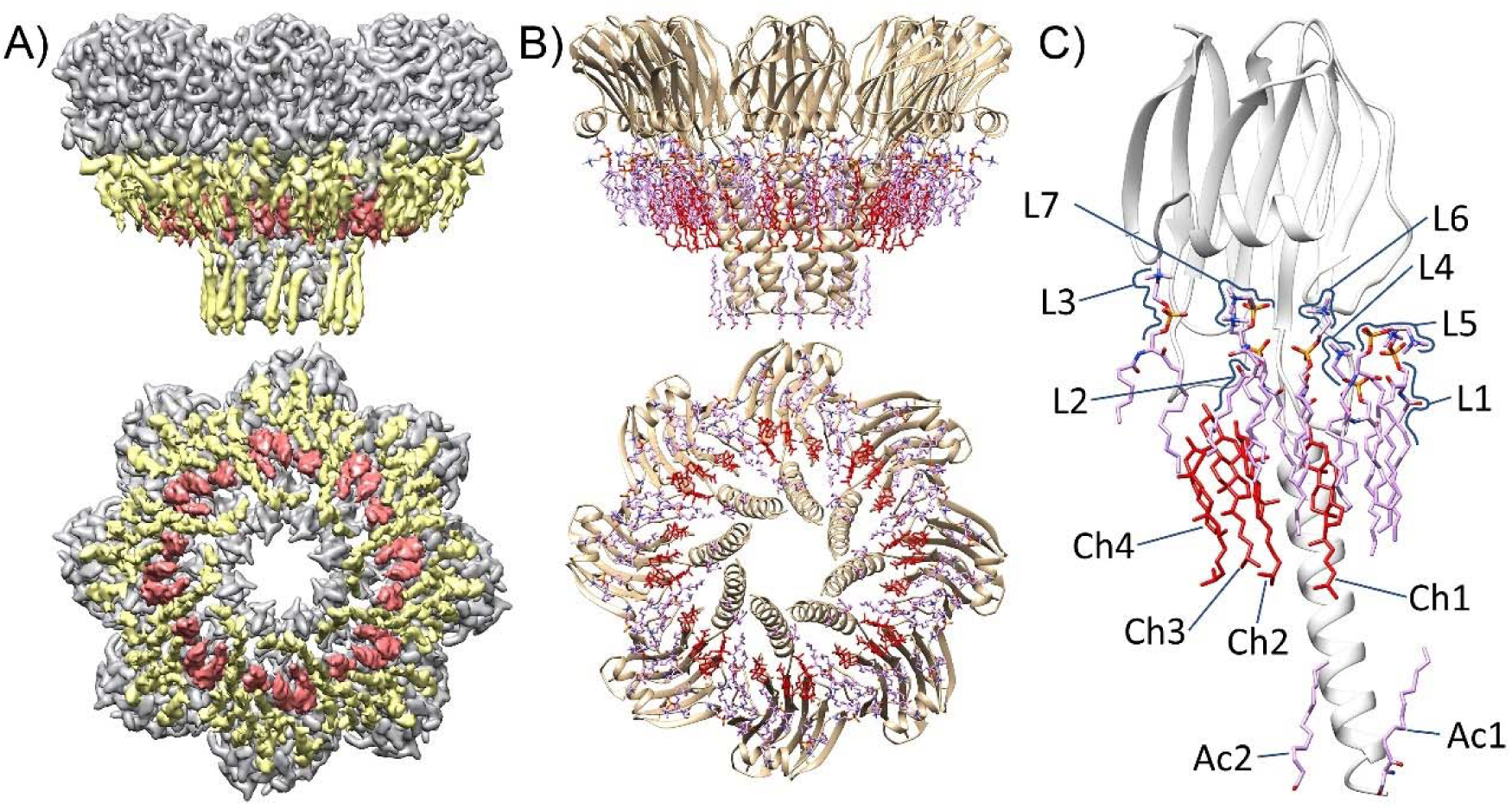
Atomic structure of FraC pore in the membrane environment. A) Side and bottom views of the CryoEM map of FraC obtained from LUVs. Densities corresponding to protein, SM and cholesterol are shown in grey, yellow and red, respectively. B) Side and bottom views of the atomic structure of the octamer with the eighty-eight lipid molecules and the sixteen acyl chains, SM are colored in pink and cholesterol in red. C) Lipid arrangement in a monomer. SM molecules are numbered from L1 to L7, cholesterol from Ch1 to Ch4, and acyl chains Ac1 to Ac2. The FraC monomer has been colored in grey for clarity.

**Figure 2.**
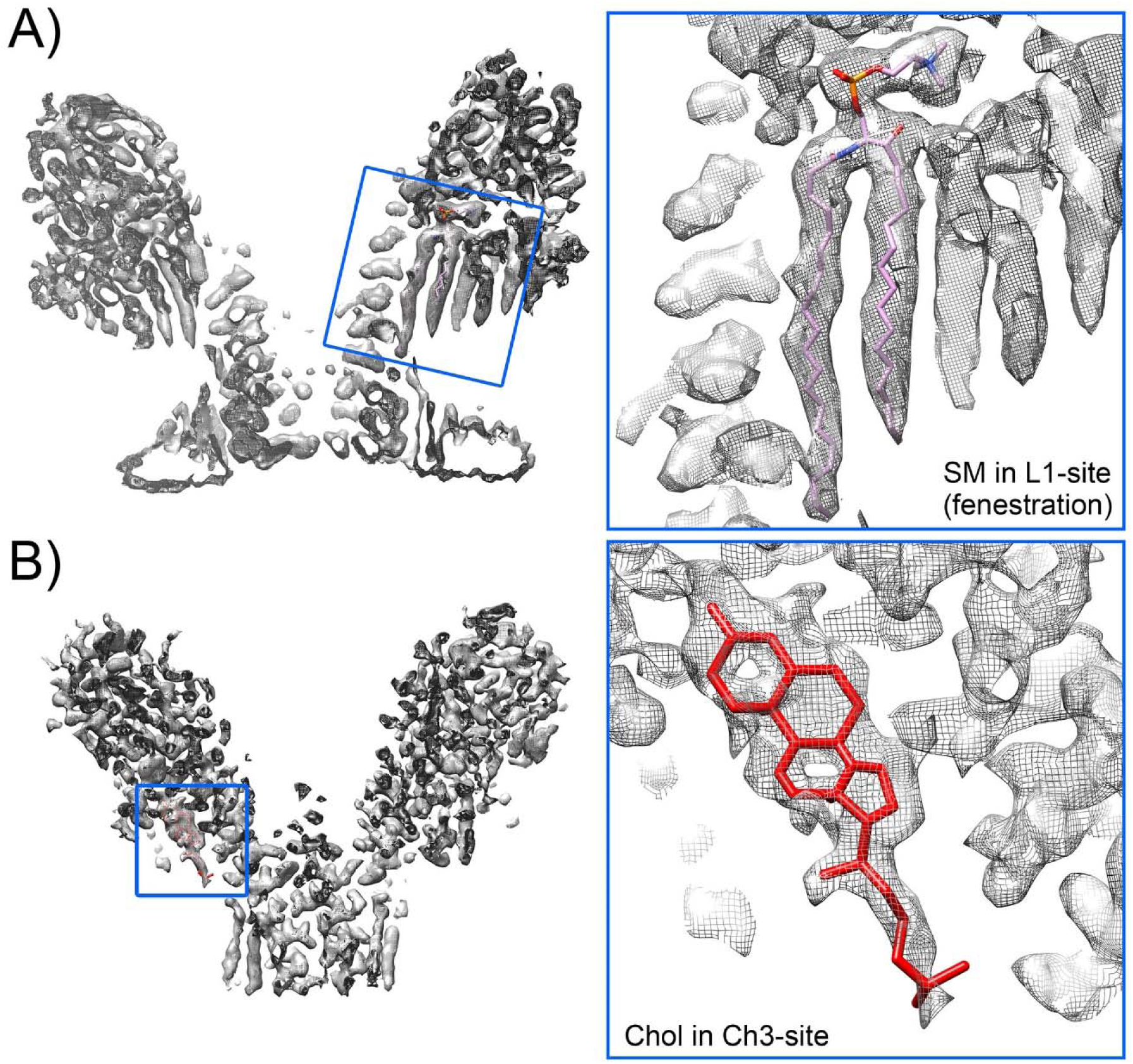
Lipid densities in FraC pores in LUVs. Two sections of the FraC map showing the density corresponding to de SM present in the fenestration marked as lipid L1 in Figure 1C (A), and the Chol marked as Ch3 in figure 1C (B).

The structure of FraC pores resolved in lipid nanodiscs reveals two stoichiometries, a majority one with eight monomers per pore and another, representing approximately 2% of the total, with a heptameric arrangement (Extended Data Fig. 4A). The octameric species, solved at 2.5 Å resolution (Extended Data Fig. 4B), exhibits a structure identical to that found in liposomes (Extended Data Fig. 4C). Nevertheless, the amount of lipids ordered around the pore is smaller, possibly due to the proximity of the scaffolding protein of the nanodisc, which sterically limits the extent of lipid molecules surrounding the pore (Extended Data Fig. 5).

The structure of the StnII pores in liposomes was determined at 2.2 Å resolution, revealing a similar arrangement to FraC and confirming the high structural similarity between the two proteins (Cα RMSD = 0.455 Å) (Extended Data Fig. 6 and 7). In the upper part of the transmembrane region, four SM and four Chol molecules per monomer are present, positioned similarly to the FraC pore. Also similar to the FraC case, the structure of StnII in lipid nanodiscs exhibits two populations, one with eight subunits and the other with seven subunits, approximately in the same proportion (Data not shown).

The comparison of the structures of FraC and StnII solved in liposomes reveals a high degree of structural homology, especially in the topological characteristics of the pore channel and some functionally important amino acids. For instance, tryptophan residues, identified through mutational studies as playing a crucial role in membrane recognition and pore formation [28], exhibit a high spatial coincidence (Extended Data Fig. 8). The most significant differences arise from the electrostatic potential on the surface of both complexes (Figure 3). FraC shows negative charges along the entire transmembrane helix of the pore produced by the amino acids Asp 3, 10 and 17, and Glu24. In contrast, StnII only has a negative region formed by the amino acids Glu 22 and 23 and Asp 18 clustered in the upper third of the transmembrane helix (Figure 3A). Moreover, FraC displays a region with distinctly positive character in the β-sandwich connection region to helix 1, contributed by amino acids Lys 30 and 32, Arg 31 and 79 and His 169. In comparison, StnII has a smaller positive charge due to the presence of only two amino acids, Lys 30 and 168. Finally, another notable difference is found at the exit of the pore on the cytoplasmic side, resulting from the sequence divergence in the first 4 amino acids at the N-terminal end. The presence of the Asp residue 3 in FraC, not resolved in the X-ray structure, causes a narrowing at the pore exit and introduces a negative charge absent in the StnII structure (Figure 3B).

**Figure 3.**
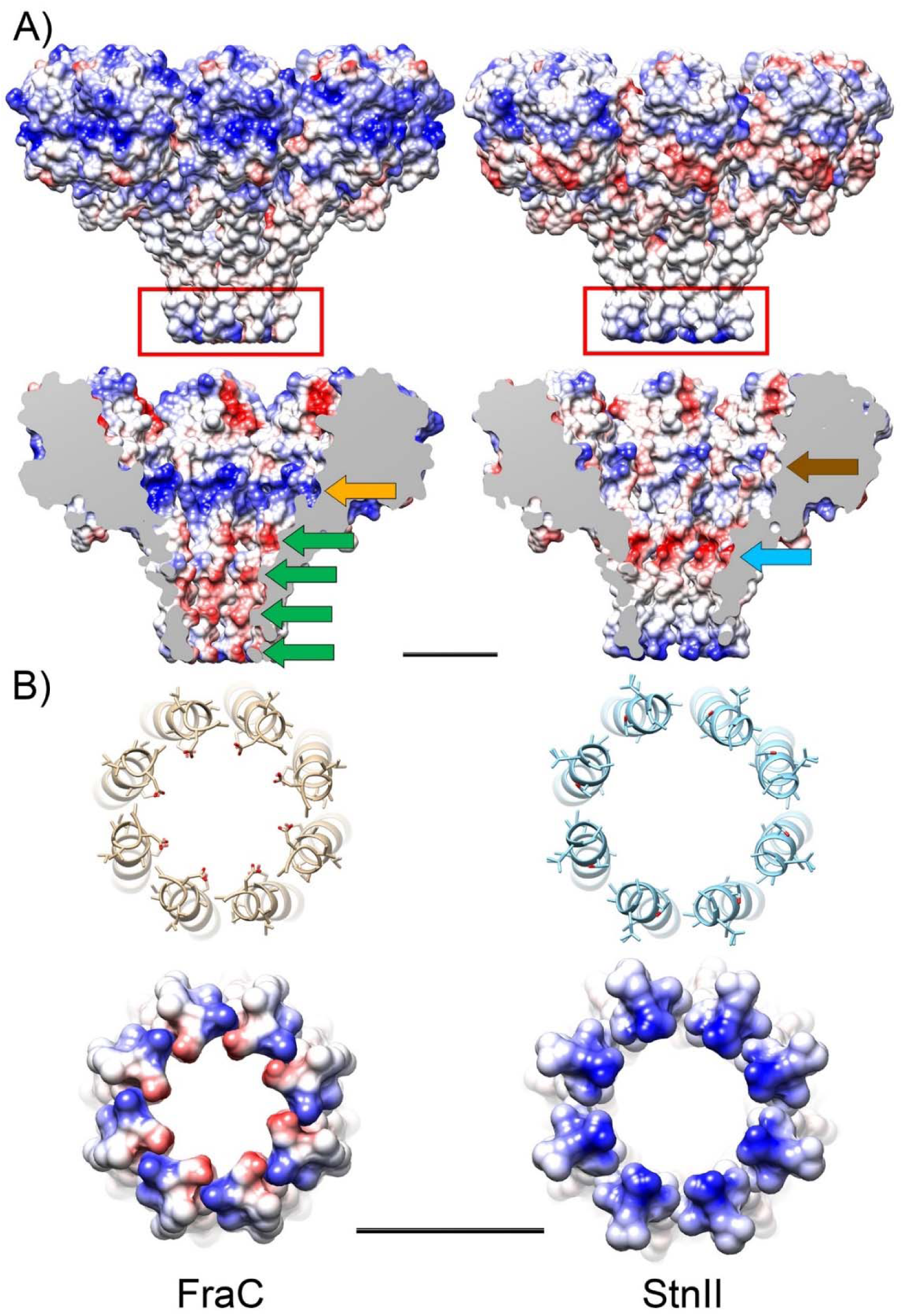
Comparison of the pore structure of FraC (left) and StnII (right). A) Electrostatic potential of the external surface (top) and inner channel of the pore (bottom). In FraC, the transmembrane helices facing the lumen of the pore are negatively charged along their entire length (green arrows), while StnII exhibits a ring at the top of the helix (blue arrow). Additionally, FraC features a ring with an intense positive charge (orange arrow), which is weaker in StnII (brown arrow). B) Detail of the region marked with a red rectangle in A, corresponding to the lower exit of the pore. The presence of the Asp 3 in FraC reduces the pore diameter and changes the charge with respect to that of StnII. Scale bar represents 25 Å).

Regarding pore stoichiometry, a number of structural and biophysical studies on these PFTs have proposed tetramers [50], octamers [13], or nonamers [51] as biological complex. In this study, we also demonstrate the presence of heptameric structures in lipid nanodiscs. Recent results [52] have led to a consensus that an arrangement of eight subunits is the most thermodynamically stable structure. Accordingly, other stoichiometries would be transient oligomers during pore assembly within the membrane [13, 53–55]. The results of this study confirm that the majority of the pores present in the membrane are octamers, with the heptamer being a structure that, although thermodynamically possible, is likely obtained due to the spatial constrictions imposed by the nanodiscs during the pore formation process.

In summary, our results demonstrate the high similarity between FraC and StnII pores, suggesting that these structural characteristics are likely shared among all sea anemone actinoporins that possess the water-soluble structure of a β-sandwich flanked by two α-helices [47]. Consequently, these all proteins would also share a common mechanism in pore formation to exert their cytolytic action.

### Role of lipids in pore structure

All structures resolved in this study, whether FraC or StnII, in LUVs or nanodiscs, exhibit a remarkable organization of the lipids around the pore, with nearly identical arrangements in all the cases (Extended Data Fig. 9). This finding points out that lipids are not merely the solvating medium of the hydrophobic part of the protein, but rather a constituent part of the pore. Since the structures are essentially identical, we will only describe in detail those obtained in LUVs, as they present higher resolution compared to the structures obtained from nanodiscs. In general terms, these results are consistent with those also described for the actinoporin-like Fav pore from the coral *Orbicella faveolata* [56].

In the FraC oligomers a total of eighty-eight lipid molecules can be resolved surrounding the upper part of the transmembrane helix. Fifty-six can be unambiguously assigned to SM (Extended Data Fig. 4), suggesting a preferential interaction of this lipid with the protein compared to DOPC, and the other thirty-two are Chol molecules. Eight of the SM molecules occupy the fenestrations between the transmembrane α-helices of consecutive monomers, while the other forty-eight surround the pore forming a wide ring. The Chol molecules are also arranged in a ring around the pore, but in the central region of the membrane. Detailed analysis of lipid distribution by monomer reveals that, among the seven SM molecules associated with each monomer (Figure 1C), the innermost three match those identified in the crystallographic structure of 4TSY [13]. These molecules are the ones closely interacting with the protein, including the one located in the fenestration. The remaining four outermost molecules, absent in the crystallographic structure, primarily interact with the protein only through their polar heads (see Extended Data Fig. 9 and 10). The four Chol molecules corresponding to each monomer interact with the hydrophobic part of the transmembrane helix on one side, and with the end of the acyl chains of the SMs on the other (see Extended Data Fig. 9 and 10).

In the case of StnII pores in LUVs, thirty-two molecules of SM and other thirty-two molecules of Chol have been resolved in positions completely similar to those of FraC (Extended Data Fig. 9). The fact that in this second case fewer lipids have been built into the density map is most likely due to a slightly lower quality of the density map on the outside of the structure, rather than to real differences in their biochemical behavior.

The close interaction between the lipids and the protein in both cases likely contributes to the high thermodynamic stability exhibited by the protein [29, 52, 57](see discussion). Among these lipids, SM L1 is of special interest because it closes the fenestration gap left between the α-helices 1 of adjacent monomers on the extracellular side of the membrane as described [13]. Our density maps also show how the acyl chains are fully extended to cover the opening between the protomers (Figure 2 and Extended Data Fig. 7). On the other hand, the Chol molecules present in both structures are located in the region that approximately corresponds to the hydrophobic core of the membrane, and their positions are rigidly fixed by interaction with the protein and the acyl chains of the SMs. This is an arrangement that fits perfectly with the very well-known preference of Chol for this sphingolipid [58]. Finally, it is also important to note that the equivalence in lipid positions between FraC and StnII pores further highlights the structural similarity between the two proteins and, presumably, their action mechanisms.

### Structure of pore intermediates

Preparations of FraC and StnII on lipid nanodiscs occasionally resulted in intermediate species composed of 4, 5, or 6 monomers, forming half-ring or arc-shaped structures (Figure 4 and Extended Data Fig. 11A). The 2D averages revealed a consistent appearance among all the monomers forming the arc, except for the one at one end, which has an additional density facing the center of the pore (Figure 4A arrow). The structure of each monomer also resembled that of the monomers in the fully assembled octameric pore, except that the helix inserted in the membrane was not defined in the arcs. All these data suggested that these structures were intermediates of the pore formation process.

**Figure 4.**
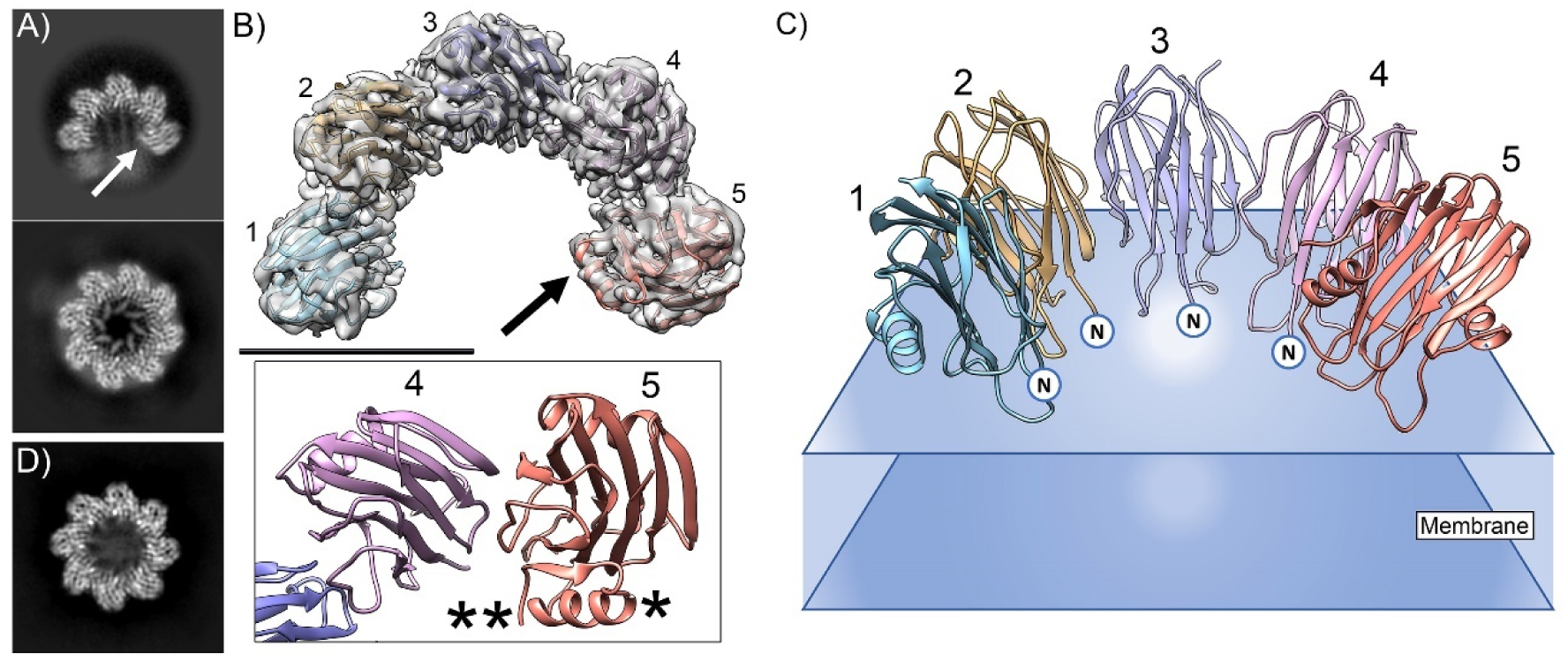
Structure of pore intermediates. A) 2D average of an intermediate pore composed of five monomers (top) compared with the 2D average of the final octameric pore (bottom). The arrow indicates the distinctive feature of the monomer at the end of the arc. B) Top: Three-dimensional reconstruction and atomic structure built in the pore intermediate of StnII. The arrow points to the α-helix 1, which folds over the β-sandwich in the last monomer and corresponds to the distinguishing feature of the 2D average. Scale Bar represents 50Å. Bottom: Detail of the atomic structure of monomer 4 and 5 showing the position of the α-helix 1 (*) and the N-terminus of the last monomer (**). C) Diagram of the atomic structure of the intermediate pore on the membrane (blue planes). The position of the unresolved extended α-helix 1 has been marked with an “N” in the first four monomers, while the last monomer is fully resolved. D) Two-dimensional average of an octameric pore in which the helices that form the pore wall are not placed in their final position, but remain extended/disordered on the surface of the membrane.

To obtain the three-dimensional reconstruction of the pore intermediates in StnII samples, CryoEM data were taken with a tilt of 30°. This approach was necessary as the nanodiscs exhibit a preferred orientation on the grid, revealing only the top views (Extended Data Fig. 11A). Since the arc structures with 5 and 6 monomers were the most abundant it was possible to attain a 3D reconstruction of these variants at resolutions of 3.5 Å and 4 Å resolution respectively (Figure 4B-C and Extended Data Fig. 11B-D). In four monomers, in the case of 5mer arc, and five monomers, for 6mer arc, it is only possible to solve the part corresponding to the β-sandwich and the α-helix 2, at the external part of the pore, but the density corresponding to α-helix 1 is not clearly defined. In the last monomer, placed at one end of the arc in both cases, it is possible to solve the complete structure and shows a folding similar to that of StnII in its soluble form, with the α-helix 1 folded over the β-sandwich (Figure 4B and Extended Data Fig. 11D). The lateral interaction between the first monomers is virtually the same as that in the complete octameric pore, while the last monomer has a slightly more closed position than would correspond to its equivalent in the complete pore sandwich (Extended Data Fig.11B and D). Furthermore, the polar heads of some lipids can be resolved in contact with the monomers and corresponds to some of those present in the complete octameric pore structure (Extended Data Fig. 11C-D).

Additionally, during 2D classification of FraC images in nanodiscs we obtained an average showing a complete octameric pore, but in which the helices forming the walls of the transmembrane part are not visible (Figure 4D). While this structure is a minority in the image set (<1%), it could suggest that the disorganization of α-helix 1 might persist even when the pore is complete, potentially representing a stage just prior to membrane piercing for final pore formation.

It is important to note that these pore intermediates were only found in nanodisc samples and not in liposomes. One of the reasons that may explain this fact is the limited membrane surface area presented by the nanodiscs to interact with the soluble protein. Consequently, there is a noticeably lower rate of monomer incorporation into the structures that lead to the formation of the final pore, resulting in a slower growth rate and, therefore, the vitrification process can isolate the intermediate structures.

### Pore formation mechanism

Based on the intermediate pore formation states obtained here and the various atomic structures of StnII, FraC and other PFTs available, it is possible to propose a detailed pore formation mechanism at the atomic level.

The atomic model built on the StnII pore intermediate structures clearly shows that in four (or five) of its monomers the α-helix 1 is deployed from β-sandwich and extended/disorganized on the membrane surface, while the monomer at the end of the arc keeps this helix folded over the protein main body. This finding points to a sequential mechanism wherein monomers are incorporated one by one over an initial monomer in the membrane, causing the arc to grow concomitantly with the helix extension process, until it eventually closes into a complete pore. The potential mechanism underlying the simultaneous incorporation of soluble monomers and the unfolding of the helix can be inferred from an examination of the different existing crystallographic structures of PFTs. Comparing these structures, two noticeable changes in the α-helix 1 and in the N-terminal region become apparent: 1) the position of the first ∼5-7 N-terminal amino acids exhibits high structural variability, presenting two conformations, one proximate to the β-sandwich and the other distanced from it (Extended Data Fig. 12A). 2) The position of Phe-14 residue (Phe 16 in FraC), conserved across all actinoporins [47], also has two main positions, one closely associated with the β-sandwich and the other displaced from its hydrophobic environment (Extended Data Fig. 12B).

Putting all these data together we can outline a mechanism for pore formation (Figure 5). In the first step, a water soluble molecule of StnII spontaneously interacts with the membrane surface [50] through its well-conserved lipid-binding pockets [11, 13, 33, 41]. This first membrane-associated monomer serves as a nucleation core for toxin oligomer growth (Figure 5A). The binding of the next monomer to the membrane, along with the ability to diffuse laterally on the membrane, enables interaction with the first monomer resulting in the formation of a dimer. During this process, the Val 57 (Val 60 in FraC) and the N-terminal end of the second monomer can displace Phe 14 of the first monomer from its hydrophobic environment by steric clash (Figure 5B-C), destabilizing the binding of α-helix 1 to the β-sandwich and causing it to unfold over the membrane. A similar effect has been described in FraC crystallographic dimers, with the displacement of the equivalent Phe 16 by Val 60 (PDB 4TSN, Extended Data Fig. 12B) [44]. This clash also causes the N-terminal end of second monomer to acquire the position separated from the main body of the protein, thus also decreasing its interaction with the β sandwich. Similarly, the binding of a third monomer will induce the unfolding of the helix of the second monomer, continuing in this manner until the pore is complete (Figure 5D-E). It is important to notice that biochemical data have shown Val 60 in FraC as a key residue involved in the oligomerization of the functional pore, and mutants substituting Phe 16 are 1000-fold less hemolytic than the wild type protein against sheep erythrocytes [44].

**Figure 5.**
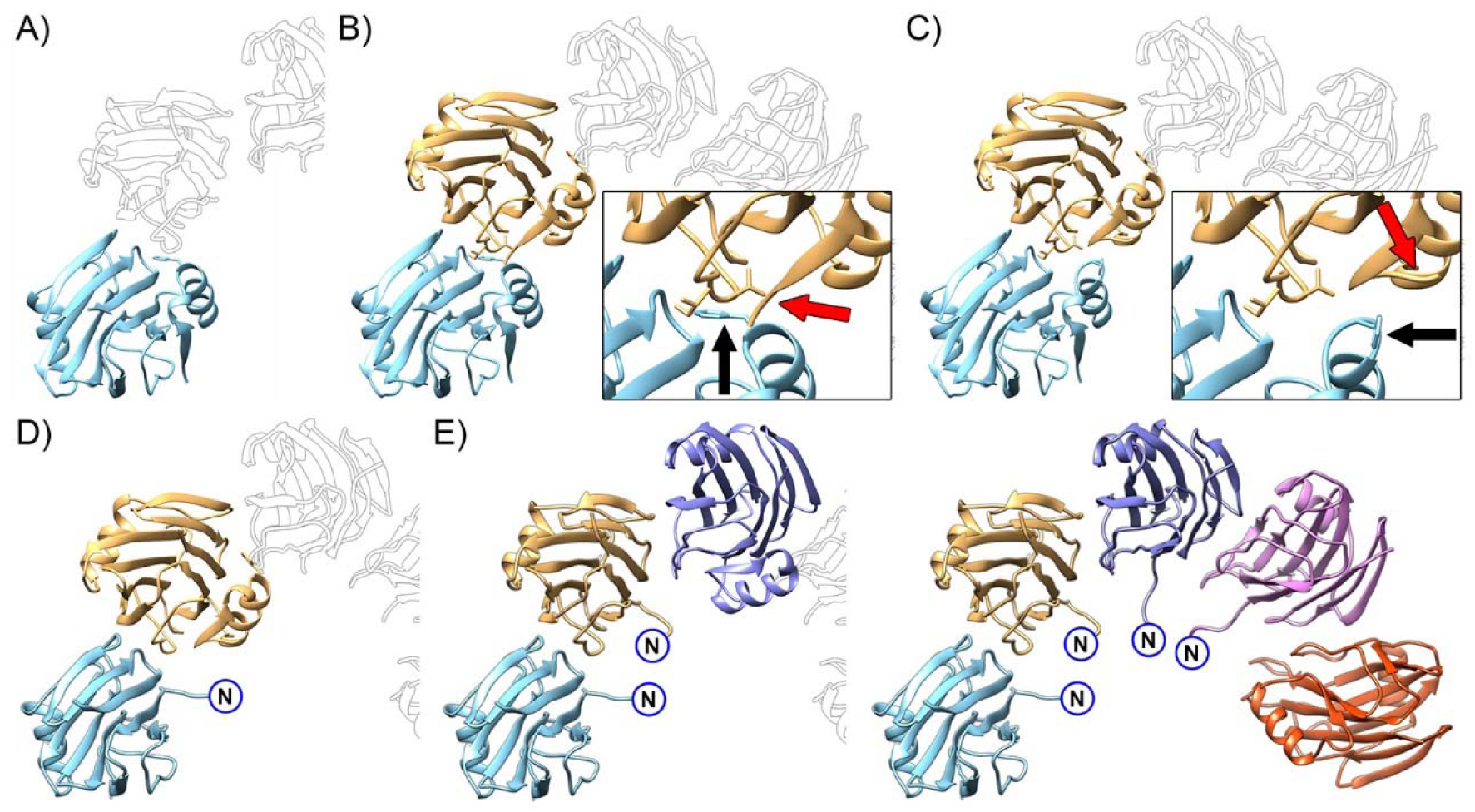
Pore formation mechanism. A) The first monomer (light blue) binds to the membrane. B) A second monomer (brown) laterally interacts and causes a clash between Phe 14 of the first monomer (black arrow) and Val 57, Ile 58 and N-terminal of the second monomer (red arrow). C) This clash results in displacement of Phe 14 from its hydrophobic pocket (black arrow) and separation of the N-terminus of the second monomer from the β-sandwich (red arrow). D) Displacement of Phe 14 destabilizes the binding of α-helix 1 to the β-sandwich and stretches it over the membrane surface. E) Successive monomer binding drives pore growth. In all panels, the position of monomers in the intermediate pore structure is outlined with black lines.

The determination of the specific moment when membrane perforation occurs cannot be precisely estimated. The images depicting pores with eight monomers, but in which the transmembrane helices are not defined (Figure 4D), might indicate the presence of a closed structure as the last step before membrane puncturing occurs and suggesting the existence of closed prepores [59]. On the other hand, studies on liposome-encapsulated substrate release have demonstrated the liberation of large molecules, exceeding the pore diameter [57]. This observation suggested that pore-forming intermediates could destabilize the membrane sufficiently to induce partial membrane rupture and lipid vesicles contents leakage [57] before the completion of the final pore. In any case, both mechanisms are not mutually exclusive and could potentially coexist during the action of the toxin.

## Discussion

In this work, we have determined the structure of two pore-forming toxins in the membrane environment using CryoEM with either LUVs or nanodiscs. Both systems have proven to be good alternatives for resolving the oligomer structure at high resolution, with LUVs yielding better resolution. In fact, to the best of our knowledge, this work presents the highest resolution of a membrane protein solved in liposomes to date. Regardless of potential experimental and technical differences in data acquisition for both samples (such as different microscopes, ice conditions, etc.), LUVs offer the advantage that pores are not subject to constrictions caused by the presence of the scaffolding protein in the nanodiscs.

One of the most striking features of the pores is the presence of a large quantity of lipids molecules and their level of organization. The resolution obtained allowed us to determine that they consist of two rings, one of SM and another of Chol. Surprisingly, no DOPC molecules were found in the rings although they constitute a majority of the lipids within the membrane composition employed. The presence of a lipid ring surrounding a pore has been shown in other cases by cryo-electron crystallography as in aquaporin-0 [60], but it could not be determined what type of lipids formed it because it was a non-specific interaction. However, our work shows a structure with two rings of specific composition and formed by a high amount, well-defined lipid molecules. Furthermore, this observation concurs with similar results described for the actinoporin-like Fav pore from the coral *O. faveolata* [56].

The general function of lipids bound to the membrane proteins can be divided into 3 types (reviewed in [61]): 1) specific binding of specific lipids acting as cofactors essential for protein function [62, 63]; 2) formation of a lipid ring (’lipid annulus’), different in composition from the bulk of the membrane [64], and which is important for the complex functionality [65]; and 3) non-selective solvation of the transmembrane domains. In actinoporins it is well established that the presence of SM is essential for their cytolytic action [21, 24–26, 31, 32, 39, 41], whereas Chol enhances efficacy but is not strictly necessary [37–40, 42]. The defined position of SMs in the structures solved here suggests that they are an essential component of the pore, such as the one found in the fenestration. However, since SM also forms a ring around the pore, this could be a case where a lipid fits into the first two types of functions mentioned. It should be noted that SM has also been shown to play a key role in membrane recognition by the soluble form of the protein, making the presence of this lipid even more critical.

Regarding Chol, a function could be to adapt/accommodate the thickness of the lipid bilayer to the length of the transmembrane fragments of the pore (extended α-helix 1). It has been shown that a discrepancy between membrane thickness and length of protein transmembrane regions can lead thermodynamically unfavorable states by potential exposure of hydrophobic zones to the solvent [66], and actually that seems to be also the case for actinoporins [34]. On the other hand, PFTs are adapted to act on the plasma membrane, which is thicker (∼45 Å) than the inner membranes of the cell (∼35-40 Å) [67]. This agrees with the values we obtained by measuring the bilayer thickness in our maps in the area closest to the pore, which are about 41-43 Å. In the solved structures, Chol accumulates in the central region of the bilayer, sterically pushing the SM outward and thereby increasing the membrane thickness to better accommodate the helices that form the pore. This hypothesis confers on Chol a role as a thermodynamic stabilizer of the pore once formed, thus confirming biochemical data indicating that it is not an essential component in the toxin action but may enhance its cytolytic performance [27, 37–40, 42]. It is also noteworthy that the composition of the plasma membrane is much richer in Chol (up to 45 mol%) and sphingolipids than those inside the cell, such as the endoplasmic reticulum (down to 5 mol%) indicating the adaptive evolution of these proteins to the plasma membrane.

The stoichiometry of FraC and StnII pores mostly corresponds to octamers, although heptamers were obtained as a minority species (∼2%) in some samples prepared on nanodiscs. The existence of these heptamers confirms the ability of these proteins to form structures with different stoichiometries, as demonstrated previously with the crystallographic structure of nonamers in FraC [51], and for other PFPs [68, 69]. Nonetheless, it seems probable that the formation of heptameric pores may be favored by constraints imposed by the nanodisc on the pore formation process.

In nanodiscs occasionally appeared arc-shaped structures that correspond to pore-forming intermediates. In those intermediates the monomer at one end retains a structure very similar to the water-soluble form of the protein, while the rest have the α-helix 1 deployed over the membrane. On the other hand, early electron microscopy studies on StnII showed its ability to form two-dimensional crystals on lipid monolayers [11, 50]. This indicates that the soluble protein can spontaneously bind on the membrane through the described sites [11, 13], and diffuse until it interacts with other monomers. Based on these data we have proposed a mechanism in which a membrane-bound monomer serves as a nucleation point for others to bind by lateral diffusion and form the pore. As described, the mechanism that drives the pore formation is the result of the lateral interaction of the monomers combined with steric clashes, resulting in the extension of the helix on the membrane surface. Finally, the timing of the puncture that perforates the membrane is not clear and may occur after the pore closes, as suggested by some microscopy data, or before the structure is complete, as indicated by certain biochemical data on membrane destabilization during pore biogenesis [57].

In summary, this study elucidates the structure of two actinoporins within the membrane environment, revealing the significant involvement of many lipids in the pore arrangement. Furthermore, we propose a mechanism of pore formation that, due to the high structural homology within this family of proteins, is likely common to all its members, significantly advancing our quantitative understanding of the mechanisms underlying pore-forming protein action. From a technical perspective, our findings demonstrate the efficacy of utilizing LUVs decorated with membrane proteins for high-resolution structure determination via cryogenic electron microscopy. Moreover, we established the utility of nanodiscs in trapping intermediates involved in dynamic processes such as pore formation. All these advances will contribute to a deeper understanding of PTFs and membrane proteins within a broader context, particularly shedding light on the nature of lipid-protein interactions.

## Experimental procedures

### Materials

DOPC, porcine brain SM, and Chol were obtained from Avanti Polar Lipids Inc. (Alabaster, AL, USA). The plasmid used to produce the nanodiscs scaffold protein (MSP1E3D1) was purchased from Addgene (Cambridge, MA, USA).

### Protein homogeneity and spectroscopic characterization

The MSP1E3D1 scaffold protein and the actinoporins FraC and StnII were produced and purified as previously described [9, 49, 70]. Homogeneity of protein samples was confirmed by 0.1% SDS-15% PAGE and amino acid analysis, both performed in standard conditions [12, 70, 71]. All protein batches were characterized prior to use by spectroscopic means which included recording their near- and far-UV circular dichroism spectra on a Jasco 715 spectropolarimeter (Easton, MD, USA) as described elsewhere [28, 29]. Proteins were dissolved at 0.2-1.0 mg/ml concentrations in 15 mM MOPS buffer, pH 7.5, containing 100 mM NaCl.

### Preparation of large unilamelar liposomes (LUV)

DOPC:SM:Chol (80:20:10) 100 nm diameter unilamellar vesicles were prepared following standard procedures routinely used in our laboratory [18, 29, 72]. Briefly, an appropriate volume of the lipids dissolved in chloroform/methanol (2:1, v/v) was dried in a Univapo 100H system (Uniequip, Martinsried, Germany) to prepare the lipid vesicles. Then, films were hydrated for 1 h at 60 °C in 20 mM Tris buffer, pH 7.4, containing 100 mM NaCl. The suspension obtained was fifteen times extruded through 0.1-μm (pore diameter) Nucleopore filters (Whatman) in a thermobarrel extruder (Lipex Biomembranes, Vancouver, Canada) at 60 °C. Vesicles were employed immediately after their preparation. Final lipid concentration was estimated from the phosphorous content of the samples [73].

### Pore assembly within LUVs

Reconstitution of the actinoporin pore (FraC or StnII) on LUVs was achieved by mixing the appropriate amounts of actinoporin (prepared in LUVs preparation buffer) and liposomes to obtain a 50:1 lipid/protein molar ratio. Incubation prior to sample analysis was conducted for at least an hour at room temperature.

### Nanodiscs reconstitution and purification

The procedure employed to reconstitute a homogeneous nanodiscs population was essentially described elsewhere [9, 49] with the particular features of the system employed. These particularities emerged from the inclusion of SM and Chol within the lipid mixtures. Lipid stocks of DOPC:SM:Chol (80:20:10), prepared at 50 mM lipid concentration in chloroform, were used to prepare the convenient lipid dried aliquots. Chloroform was evaporated under a stream of nitrogen. Then, the tubes were placed for 3 hours into a desiccator under high vacuum in order to remove all traces of the organic solvent. Buffer, containing 200 mM sodium cholate at pH 8.0, was employed to rehydrate the lipid film. Typically, cholate was added up to two-fold the lipid concentration used. Tubes were then vigorously vortexed, heated at 60°C and sonicated. Then the MSP1E3D1 scaffold protein was added to the lipid:detergent mixture to a final protein/lipid ratio of approximately 1/100 and incubated at room temperature for 15 minutes. Following, detergent was removed after extensive dialysis against 20 mM Tris-HCl, pH 7.4, containing 100 mM NaCl and 0.5 mM EDTA (the chelating agent only during the first dialysis treatment). The final dialyzed solution was centrifuged, and the clear supernatant loaded onto a calibrated Superdex 200 column, equilibrated in the same buffer employed for the dialysis and attached to an ÄKTA purifier FPLC system (Amersham Biosciences, UK). Th. The resulting chromatogram yielded a symmetrical peak corresponding to the expected molecular size, which was then pooled and labeled as purified empty nanodiscs.

### Pore assembly within nanodiscs

Pore assembly was performed with the aim of mimicking the real situation encountered by actinoporins [9]. Thus, instead of reconstituting the pore simultaneously with the assembly of the nanodiscs particles, the toxic protein was added to the purified empty nanodiscs preparation at a StnII or FraC/MSP1E3D1 molar ratio of 10/1. This mixture was incubated at room temperature for 1-2 hours and then employed to prepare the grids used for the cryoEM analysis. The buffer employed was also 20 mM Tris-HCl, pH 7.4, containing 100 mM NaCl. The use of the chromatographically purified nanodiscs faction provided the additional advantage that the possible influence of the presence of traces of detergent could be completely ruled out.

### Cryo-EM microscopy

Grid preparation and data acquisition of LUVs containing Frac or StnII. Samples were diluted 5-fold and stained for 1 minute with 2% uranyl acetate. The samples were examined using a JEOL 1400 Flash microscope (JEOL, Tokyo, Japan) to verify concentration and quality. 3 µL of undiluted proteoliposome sample was applied to glow-discharged Quantifoil Cu/Rh R 2/2 Cu 200 mesh grids and blotted away for 3 s and plunge-frozen in liquid ethane. Resulting grids were screened on Talos Arctica microscope (Thermo Scientific) operated at 200 kV. Selected grids were used for high resolution imaging on a 300 kV Titan Krios G4 (Thermo Scientific) instrument equipped with cold field emission electron gun (E-CFEG, Thermo Scientific), Selectris X energy filter (Thermo Scientific) and Falcon 4i direct electron detector (Thermo Scientific) using a pixel size of 0.73 Å/pix and a total dose of 52 e–/Å2. 14,002 and 9,088 movies in EER file format [74] were recorded for StnII and FraC samples, respectively.

Grid preparation and data acquisition of nanodiscs containing Frac or StnII. The concentration of nanodiscs prepared as described above was initially assessed using negative staining. Samples were diluted 10- to 100-fold and screened using negative stain as described in the section above. Dilutions showing a compact and uniform distribution of nanodiscs on the grid surface were selected for cryoEM analysis. CryoEM grids were prepared by depositing 3 μl of the selected concentration onto glow-discharged Quantifoil Cu/Rh R 2/2 grids, followed by vitrification using a Vitrobot Mark IV (Thermo Fisher). The grid quality was assessed using a Talos Arctica microscope (Thermo Fisher) operated at 200 kV. The optimal grids were chosen for subsequent data acquisition on a Titan Krios G3i microscope (Thermo Fisher) at Diamond Light Source(Oxfordshire, UK) using a K3 (Gatan) direct detector. StnII grids displaying higher proportion of half-ring structures were imaged on a 300 kV Titan Krios G4 (Thermo Scientific) instrument equipped with E-CFEG (Thermo Scientific), Selectris X energy filter (Thermo Scientific) and Falcon 4i direct electron detector (Thermo Scientific). Pixel size of 0.546 Å/pix and a total dose of 45 e–/Å2 were used to collect 13,427 movies in EER file format.

### Image processing

#### FraC and StnII pores on LUVs

FraC and StnII pores embedded in LUVs were processed using cryoSPARC [75]. In short, movies were rendered on a 4k x 4k grid, motion corrected and dose-weighted using Patch Motion Correction job type. Next, the contrast transfer function (CTF) parameters were determined using Patch CTF job type. Around 10% of micrographs were removed based on their estimated resolution and drift quality parameters. The selected images were subjected to Topaz [76] and template-based particle picking. 2.9M and 3.8M particles were picked for FraC and StnII datasets, respectively. Two rounds of 2D classification were performed and 2D classes showing clear actinoporin features were selected. The resulting particle sets underwent multiple rounds of Ab initio, Heterogeneous Refinement and 2D classification jobs. This process led to selection of a subset of 221k (FraC) and 180k particles (StnII) which were both refined to a 2.1 Å resolution EM density maps using Non-uniform Refinement job type and C8-fold symmetry.

#### FraC pores on lipid nanodiscs

All the image processing was carried out inside Scipion3 wrapper [77]. For FraC 23.465 movies were collected at a calibrated sampling ratio of 1.06 Å/pix. Frames were aligned using MotionCor2 software [78] and CTF was calculated using CTFfind4 [79]. Over 22.1 million of particles were picked and extracted using Xmipp software [80] and 2D classified using Cryosprac software [75]. This procedure yielded approximately 5 million high quality particles, with around 95% being top views and the rest being side and tilted views. At this point, top views showing C7 symmetry (∼2%) were discarded. To avoid overrepresentation of the top views in the spatial sampling during the 3D reconstruction, around 5% of these particles were selected following an image resolution criterion in the different defocus groups of micrographs. All side and tilted views, and the selected top views were used for further 3D classification and reconstruction (∼493K particles). The initial model was generated using Cryosprac [75] or Relion4 [81, 82] showing similar results. For 3D classification the selected particles were rescaled at 1.5 Å/pix and 128x128 pixels and classified in 3 classes applying C8 symmetry using Relion 4 software. The particles of the class that showed better resolution (123K) were selected and re-extracted from original micrographs at 1.06 Å/pix and 300x300 pixels and used for the final reconstruction using Cryosparc Non-Uniform Refinement procedure [75] applying C8 symmetry.

*Visualization and figures* were created using Chimera software [83].

#### StnII intermediate pore on lipid nanodiscs

StnII intermediate pore movies were processed using the same general processing strategy as described for LUV-embedded FraC and StnII particles. Briefly, movies were motion and CTF corrected and curated based on their quality. The resulting 10,085 micrographs were used for template-based particle picking. 5M particles were picked and extensively 2D-classified yielding 429k particles which were further classified using Ab initio and Heterogeneous Refinement job types. The final subsets of 272k (5-mer) and 52k (6-mer) particles were refined to respective 3.1 Å and 4.0 Å resolutions using C1 symmetry and Non-uniform Refinement job type.

#### Atomic Structures building

A single monomer from Fra C atomic structure (PDB 4TSY) was rigidly fitted into the corresponding density map using Chimera [83]. The model was then subjected to a double real-space refinement, initially manually using COOT [84], and subsequently using an automatic procedure with PHENIX version 1.21-5207-000 [85]. The restraints employed in the real-space refinement included both, standard (bond, angle, planarity, chirality, dihedral, and non-bonded repulsion) as well as additional restraints (Ramachandran plot, C-beta deviations, rotamer, and secondary structure). A local grid search-based fit was included in the refinement strategy to resolve side-chain outliers (rotamers or poor map regions fitting). Following several rounds of real-space refinement, a stable final model was obtained and validated using the phenix_validation_cryoem module in PHENIX. Finally, the octameric model was generated using the Chimera symmetry command and further refined in PHENIX. The same procedure was followed for the complete pore structure of Stn II. The half-pore atomic structure for Stn II was generated de novo manually, with all molecules in the map constructed and refined in PHENIX.

## Accession codes

CryoEM maps and atomic coordinates have been deposited in the Electron Microscopy Data Bank (EMDB) and the Protein Data Bank (PDB) under the following (provisional) accession codes: D_1292139812 for FraC in liposomes, D_1292139813 for FraC in nanodiscs, D_1292139815 for StnII in liposomes, D_1292139816 for the five-monomer StnII half pore, and D_1292139817 for the six-monomer StnII half pore. *Deposition in progress*

## Acknowledgments

This work was supported by grant PID2020-117752RB-I00 financed by MCIU/AEI/10.13039/501100011033 and FEDER, UE and grant TED2021-132748B-I00 financed by the European Union “NextGeneration EU”/PRTR (to J.M.-B.). UCM-Banco Santander Grants PR87/19-22556 and PR108/20-26896 and UnaEuropa (Unano) SF2106 (to A.M.-P.). UCM-Banco Santander Grant PR3/23-30816 (to S.G.-L.). ERT is supported by a UCM-Banco Santander PhD fellowship grant and the European Research Council (ERC) under the European Union’s Horizon 2020 research and innovation program [850974]. J.P.-O. is recipient of a doctoral fellowship from ISB/ÅA and a postdoctoral grant from the Magnus Ehrnrooth Foundation. D.H.-M. thanks Complutense University of Madrid and Banco Santander for a PhD fellowship (CT82/20-CT83/20). CNB-CSIC acknowledges support from the Severo Ochoa Program for Centers of Excellence in R&D (CEX2023-001386-S).

The authors would like to thank Diamond for access and support of the cryoEM facilities at the UK national electron Bio-Imaging Centre (eBIC), proposals BI22006 and BI30374. To Instruct-ERIC for facilitate access for cryoEM screening through the project PID: 18916 and to the cryoEM facility of the Centro Nacional de Biotecnología in Madrid (CNB-CSIC).

## Contributions

Experiments were designed and conceived by R.A., C.S., S.M., J.G.G., A.M.P., S.G.L. and J.M.B. Protein production and purification and pore preparation experiments were performed by E.R.T., D.H.M. and S.G.L. MSP for nanodiscs was prepared by E.R.T. and S.G.L. Cryo-EM structural experiments and analyses were performed by R.A., C.S., S.M., J.P.O., D.C., E.A.P. and J.M.B. J.G.G., A.M.P., S.G.L. and J.M.B. supervised the work. A.M.P., S.G.L. and J.M.B. wrote the manuscript. All of the authors contributed to editing the manuscript, discussing the results and supporting its conclusions.

## Competing interests

The authors declare no competing interests.

## EXTENDED DATA

**Extended Data Figure 1.**
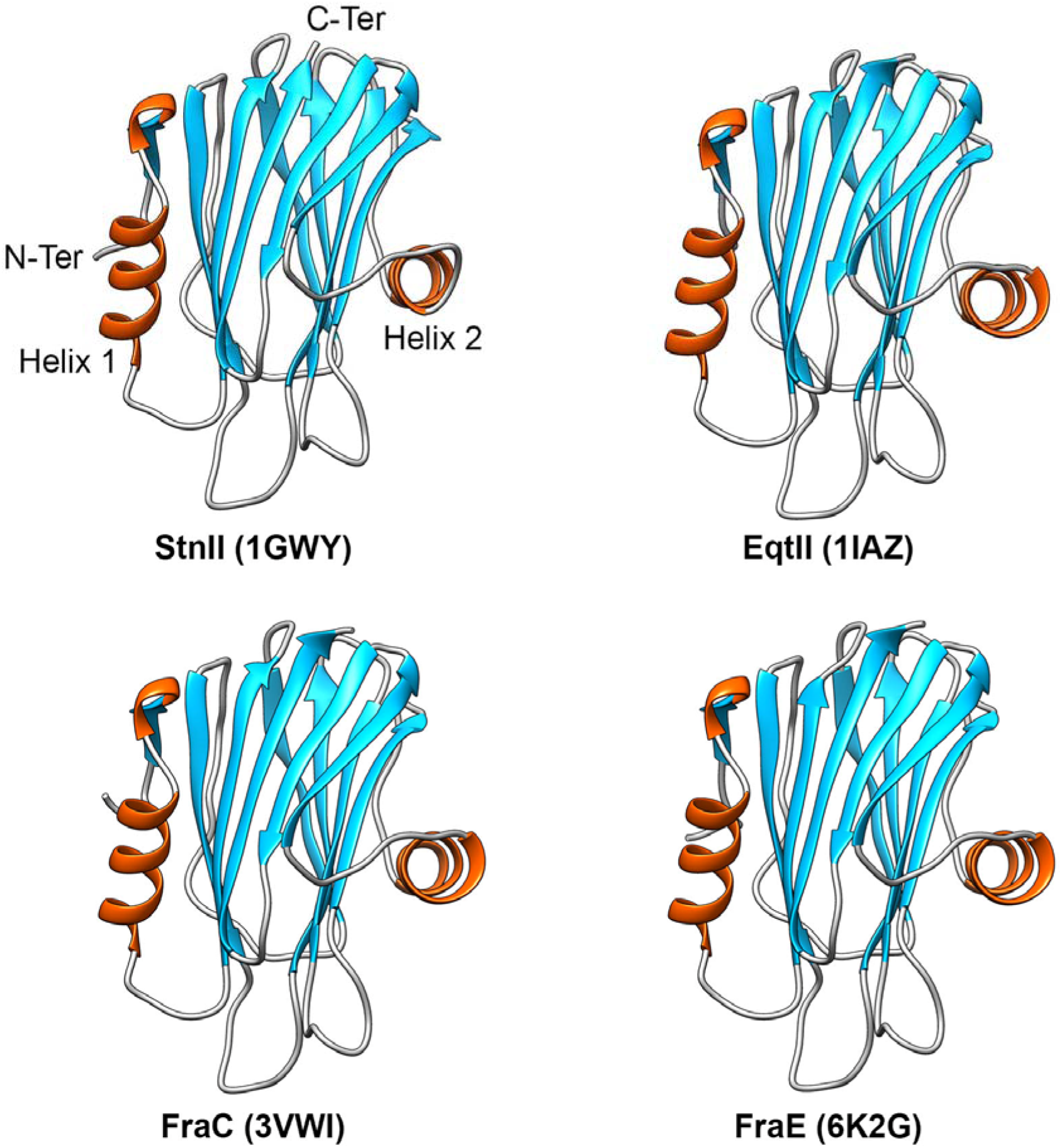
Three-dimensional structure of soluble form of best known actinoporins. Sticholysin II from *Stichodactyla helianthus* (PDB 1GWY, [11]), equinatoxin II from *Actinia equina* (PDB 1IAZ, [10]), fragaceatoxin C (PDB 3VWI, [13]) and E (PDB 6K2G, [13]) from *Actinia fragacea*. The α helices are depicted in red, β-strands are in light blue and loops in light grey. The structure primarily is composed of two α-helices flanking a β-sandwich. The helix at the N-terminal (α-helix 1, left side) is responsible for detaching, extending, and forming the pore walls.

**Extended Data Figure 2.**
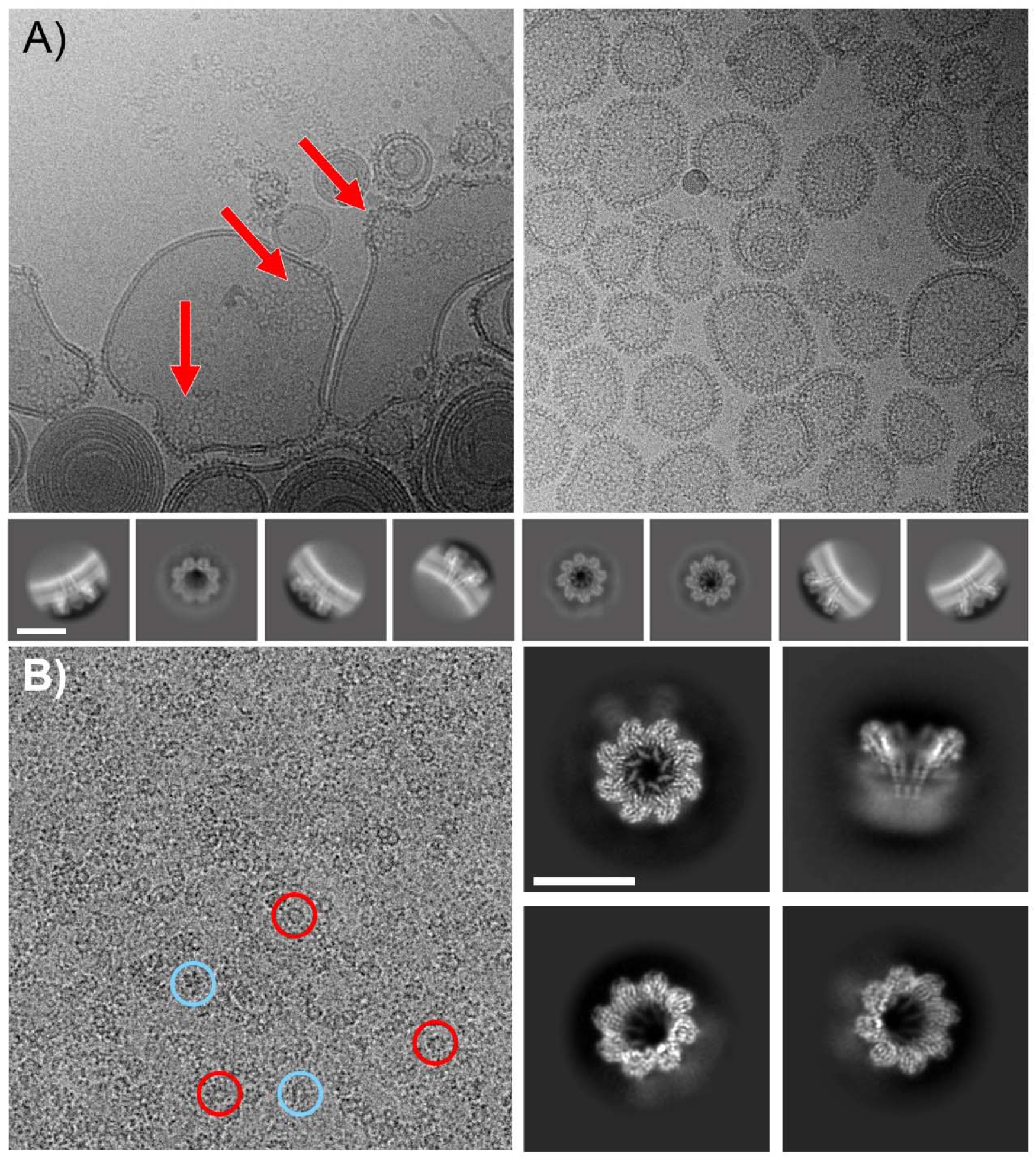
FraC pores on the membrane environment. A) FraC pores prepared on LUVs. Left: When the protein/lipid ratio is low, the formed pores are typically grouped in clusters (indicated by arrows), leaving parts of the membrane bare. These clusters tend to increase the curvature of the membrane, causing deformations in the liposome. Rigth: At higher protein/lipid ratios, the vesicles are fully decorated with pores. These samples were used to determine the pore structure in LUVs. Bottom row: Two-dimensional averages obtained by processing the pore images in LUVs. B) Left: FraC pores prepared on lipid nanodiscs. Top and side views are marked with red and blue circles respectively. Right: Two-dimensional averages of top, side and two tilted views of the nanodisc containing the pores. Nanodisc density is clearly visible in the 2D averages depicting side views. Scale Bar 100 Å.

**Extended Data Figure 3.**
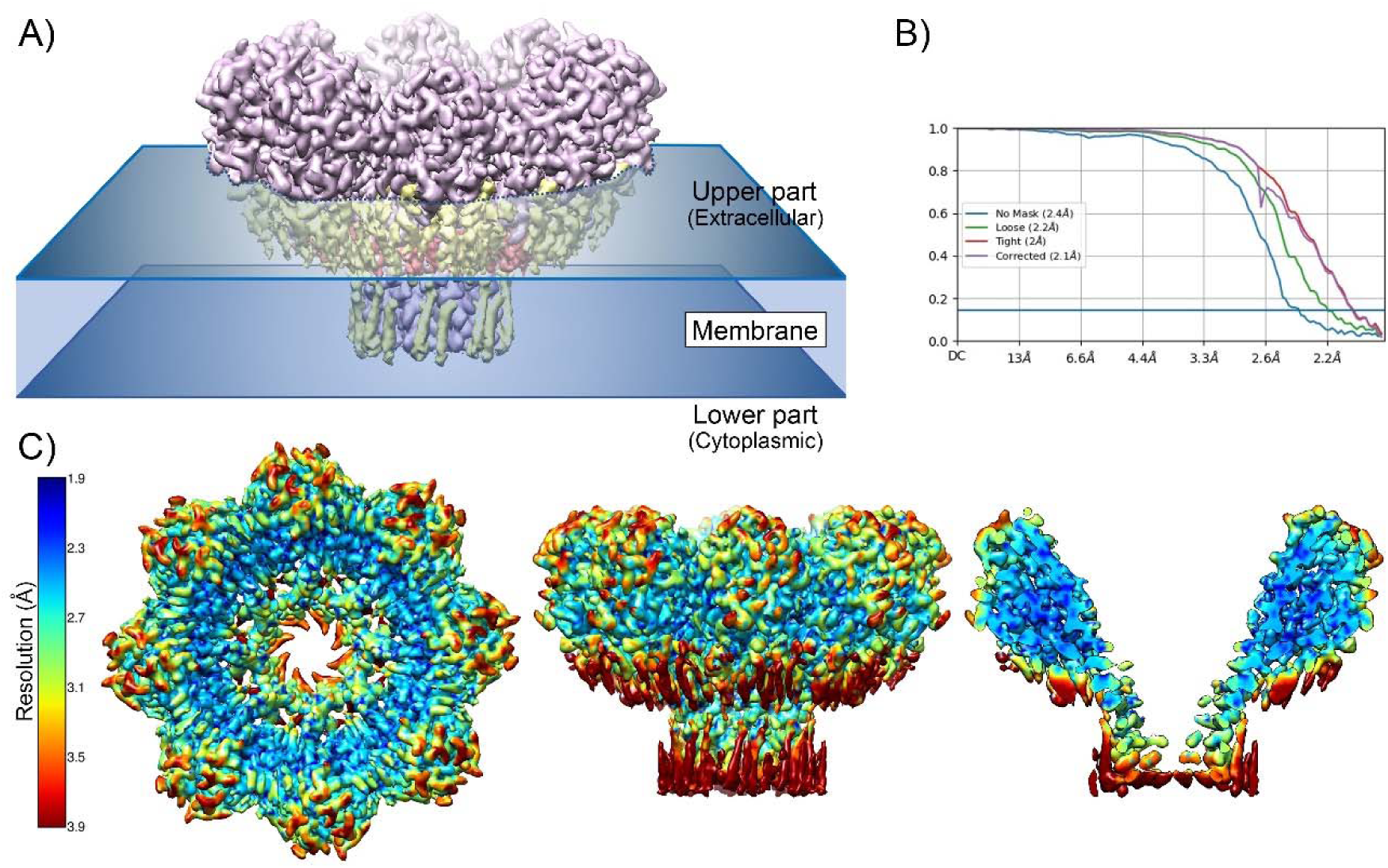
FraC pores in LUVs. A) Diagram of FraC pores in LUVs. The upper and lower part corresponds to the extracellular and cytoplasmic region respectively in the natural environment. B) Global Fourier Shell Correlation obtained with Cryosparc software [75]. C) Local resolution map calculated using MonoRes software [86]. From left to right top and side views and a central slab to show resolution of internal regions.

**Extended Data Figure 4.**
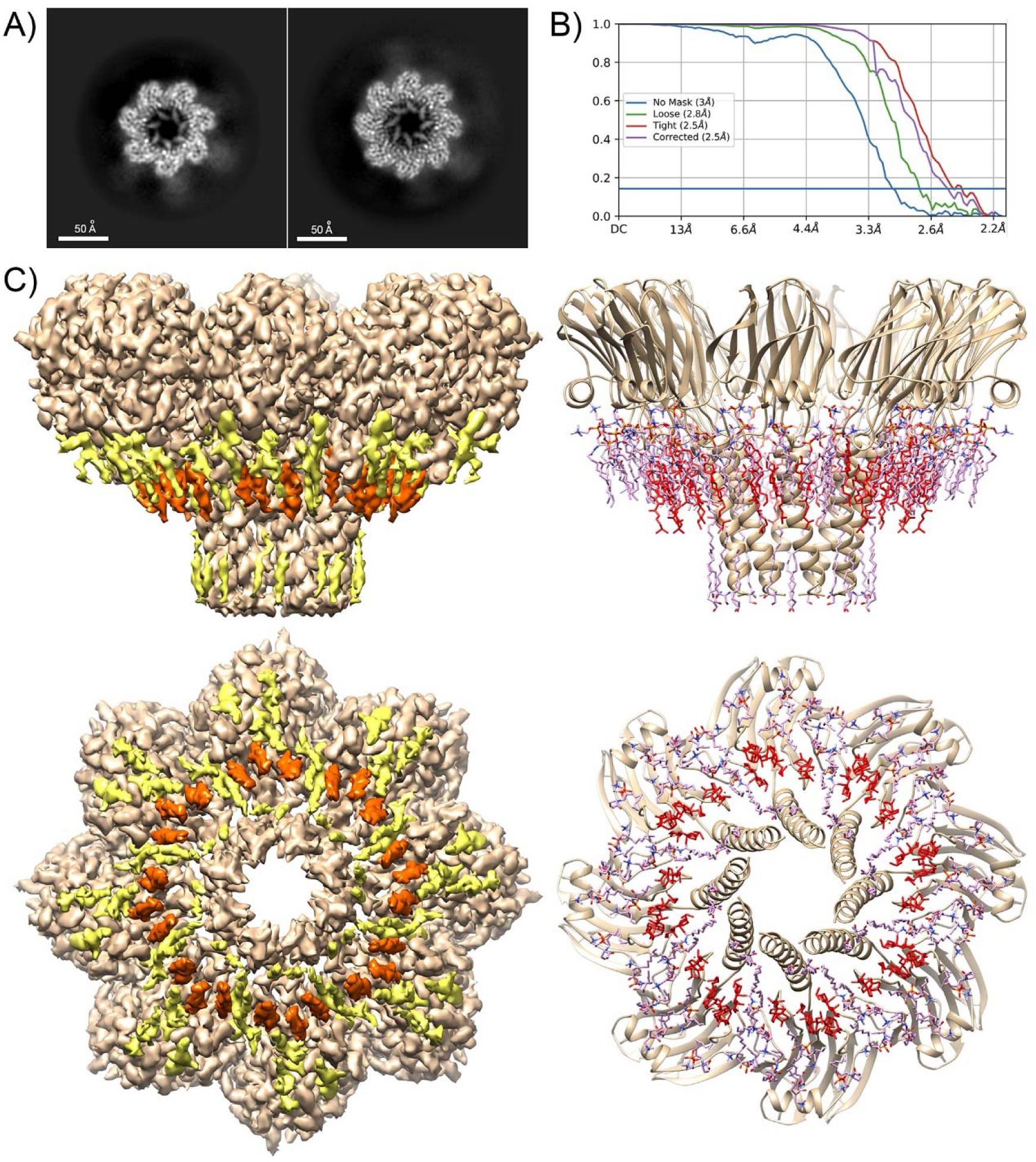
CryoEM structure of FraC in lipid nanodiscs. A) 2D average of top view showing the heptameric pore in comparison with the octameric one. B) Global resolution curve of FraC structure solved in lipid nanodiscs. C) Left: Side and bottom views of the 3D map of FraC octameric pore obtained in lipid nanodiscs. Protein is depicted in brown, SM in yellow and cholesterol in red. Right: The corresponding views of the atomic structure build in the density map.

**Extended Data Figure 5.**
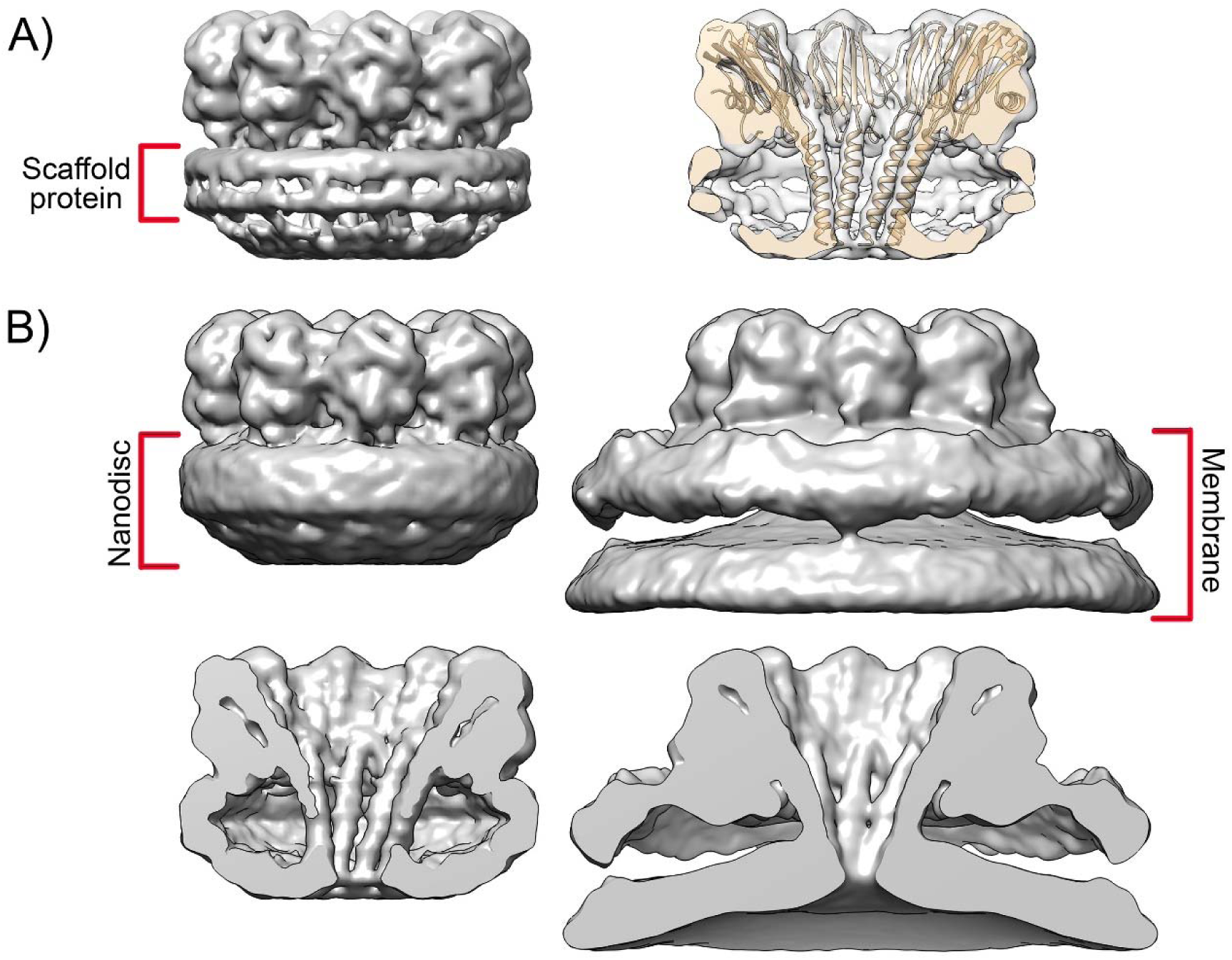
CryoEM maps of FraC in nanodiscs vs LUVs. A) Low-resolution reconstruction of FraC in nanodiscs indicating the position of the scaffold protein (left) and a cutoff revealing the docking of the atomic structure of the protein. B) Comparison of low-resolution structures of FraC in nanodiscs (left) and in LUVs (right) at ∼1.8 σ to show the extent of the nanodiscs and the membrane surrounding the pore, respectively. The proximity of the scaffold protein could represent a restriction for the arrangement of the lipids in the case of the nanodisc.

**Extended Data Figure 6.**
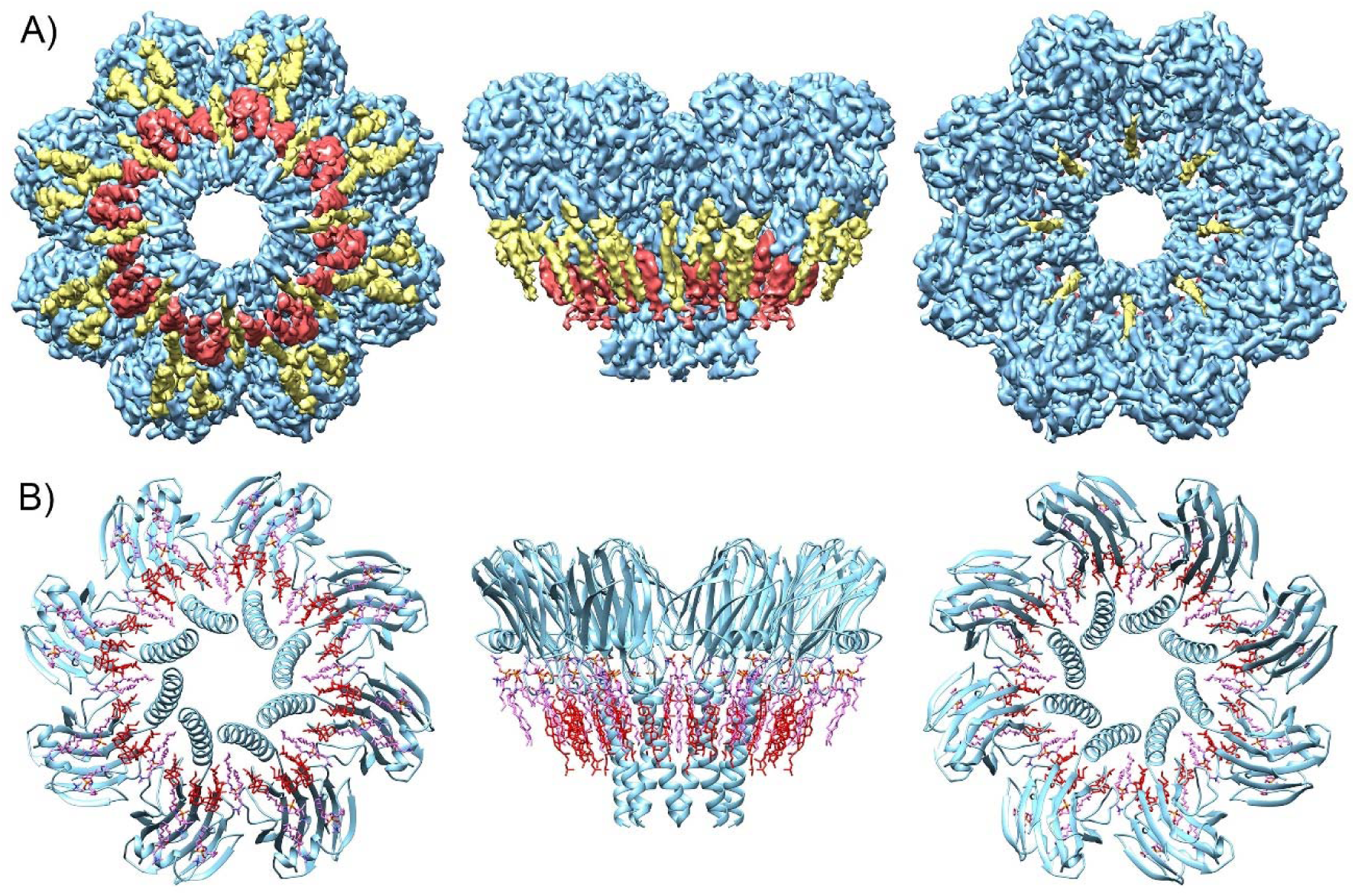
Atomic structure of StnII pore in the membrane environment. A) Bottom, side and top views of the density map obtained from LUVs preparations. Protein, SM and cholesterol are shown in light blue, yellow and red respectively. B) Atomic structure determined using the density map.

**Extended Data Figure 7.**
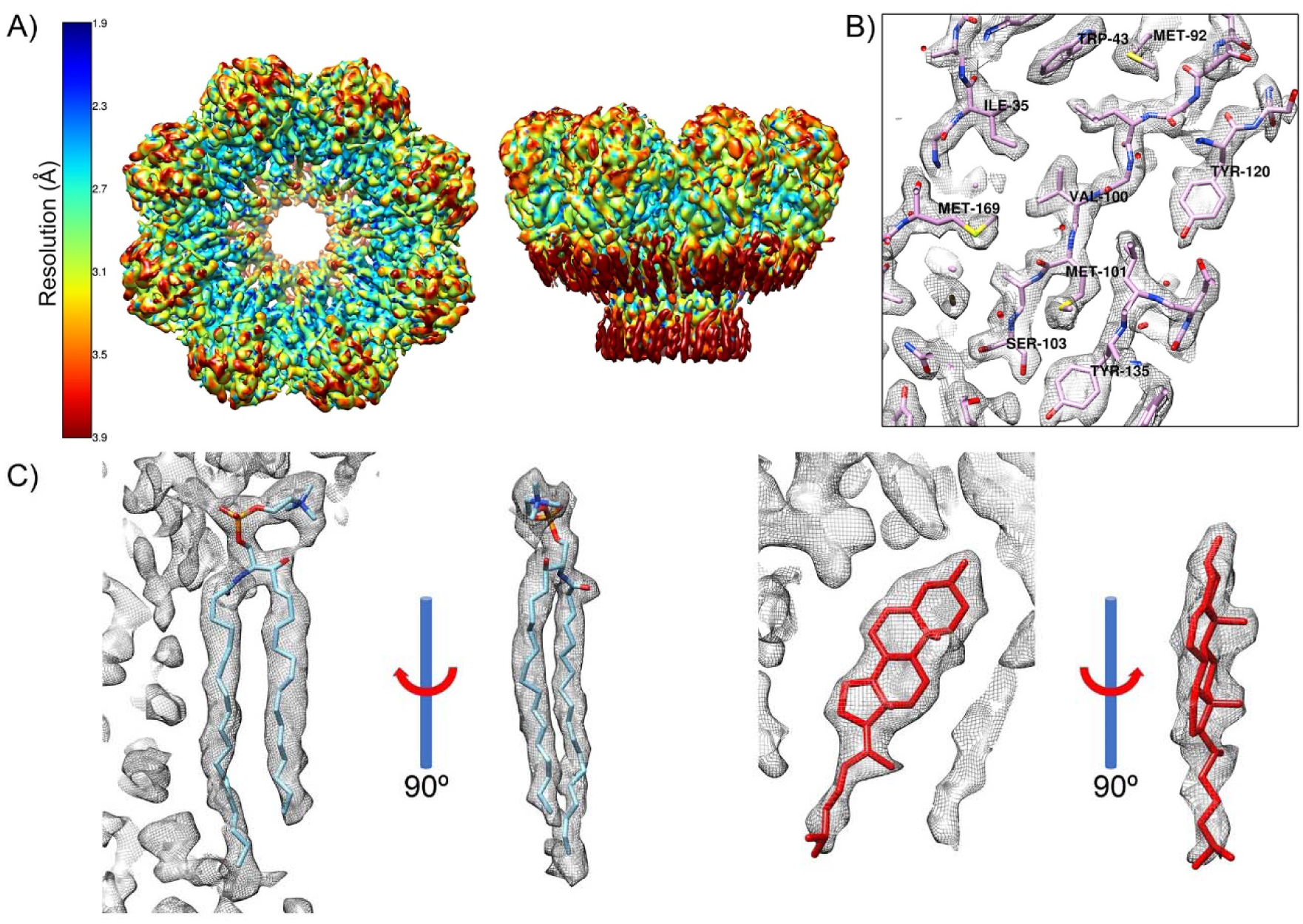
Resolution of StnII pores in LUVs. A) Top and side views of the local resolution map calculated using MonoRes software [86]. B) An example of the resolution of the density map (region around Val 100). C) Two views of the density map corresponding to the SM in the fenestration (left), and one Chol (right).

**Extended Data Figure 8.**
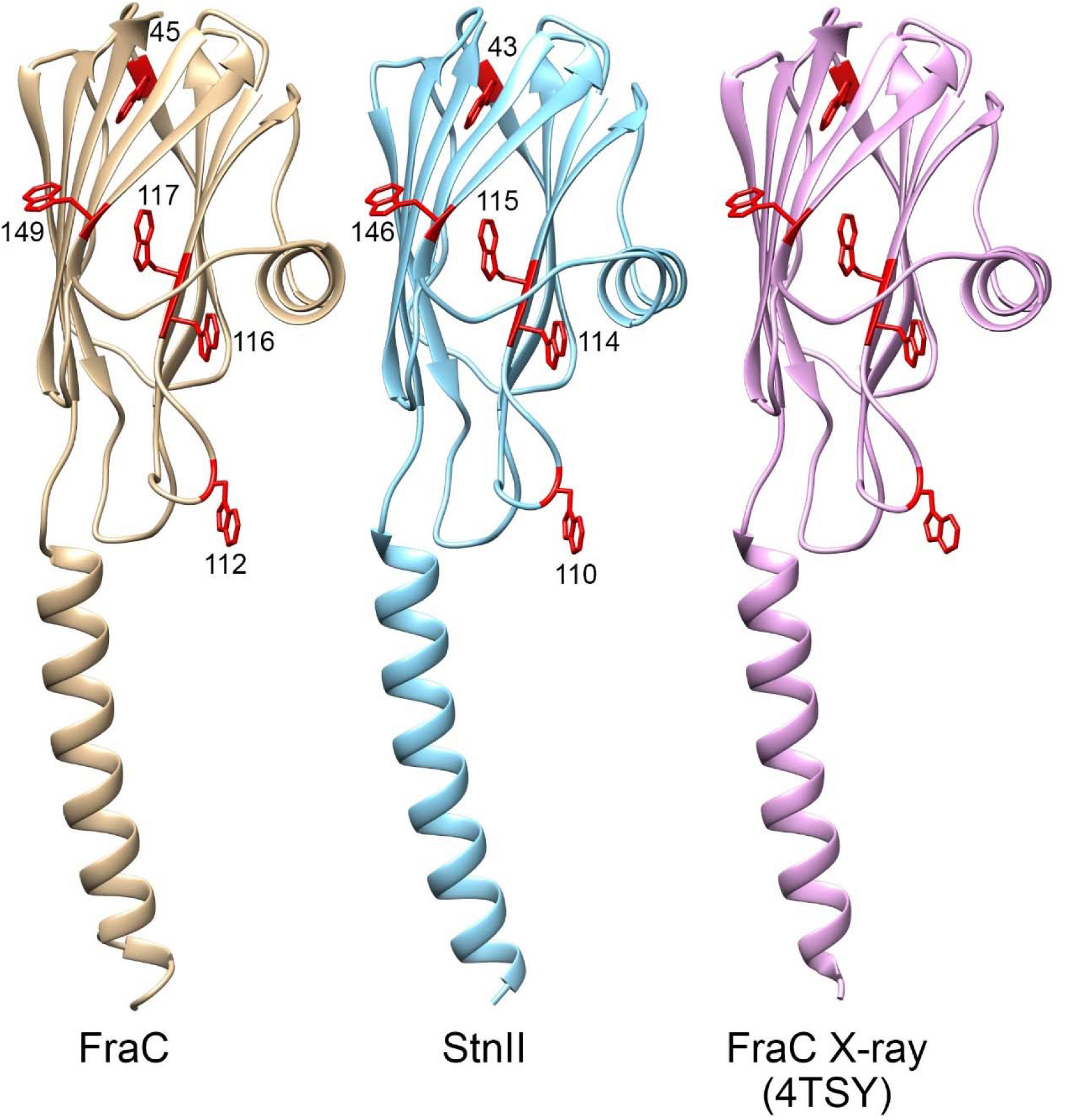
The comparison of the position of the five tryptophan residues present in the structures of FraC and StnII pores in lipid membranes by CryoEM (the two on the left), and X-ray crystallography (right, PDB 4TSY) reveals a very high spatial coincidence. The RMSD between FraC and 4TSY is 0.407 Å, 0.339 Å between FraC and StnII, and 0.471 Å between StnII and 4TSY. Tryptophan residues play a crucial role in membrane recognition, protein structure stabilization, and pore formation in anemone αPFTs and are, therefore, highly conserved. Mutagenic studies [28] have revealed that FraC W45 and W117 (StnII W43 and W115) play a determinant role in maintaining the high thermostability of the water-soluble monomer, while FraC W149 (StnII W146) provides specific interactions for protomer-protomer assembly. Finally, FraC W112 and W116 (StnII W110 and W114) sustain a hydrophobic effect, which appears to be one of the major driving forces for membrane binding. (for clarity only one monomer is shown).

**Extended Data Figure 9.**
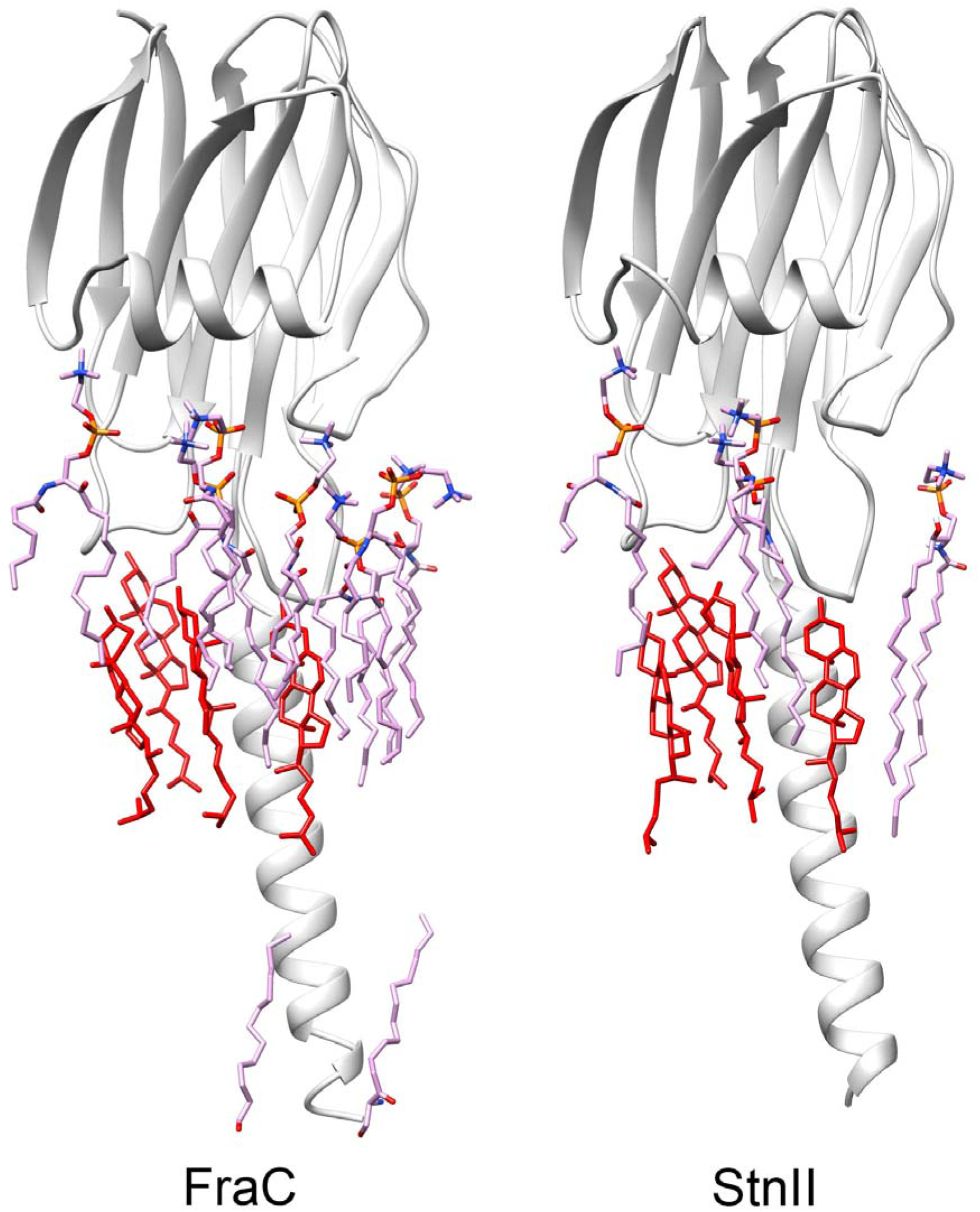
Comparison of the lipid distribution per monomer in FraC y StnII pores formed in LUVs. Left FraC and right StnII. Protein monomers are shown in grey, SM is shown in pink and Chol in red. The acyl chains at the bottom region cannot be assigned to a particular lipid type of those present in the membrane (SM or DOPC).

**Extended Data Figure 10.**
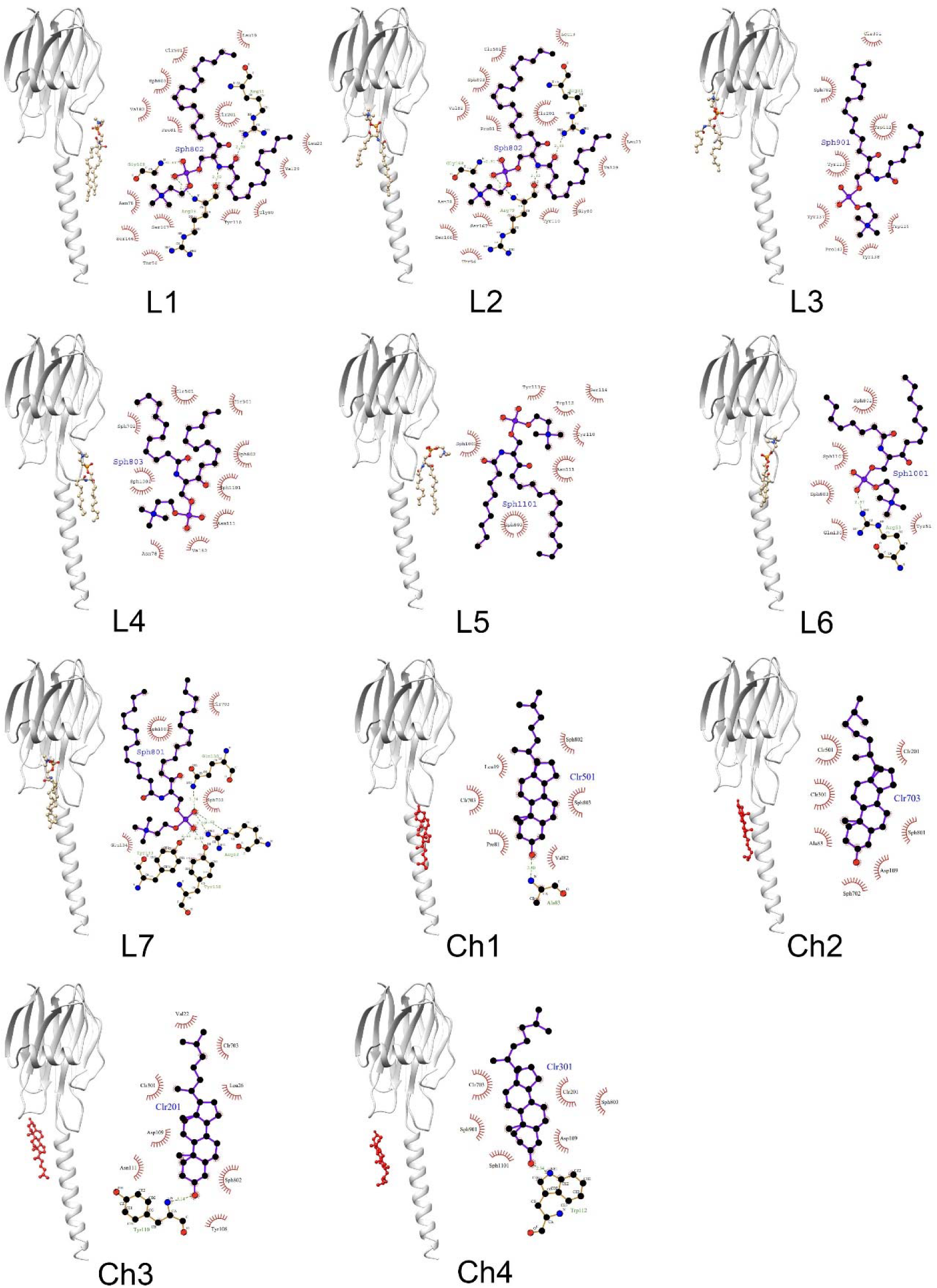
Interactions of lipids in FraC in pores formed in liposomes. SM molecules have been named for L1 to L7 and Chol from Ch1 to Ch4. Diagrams have been calculated with LigPlot+ software [87].

**Extended Data Figure 11.**
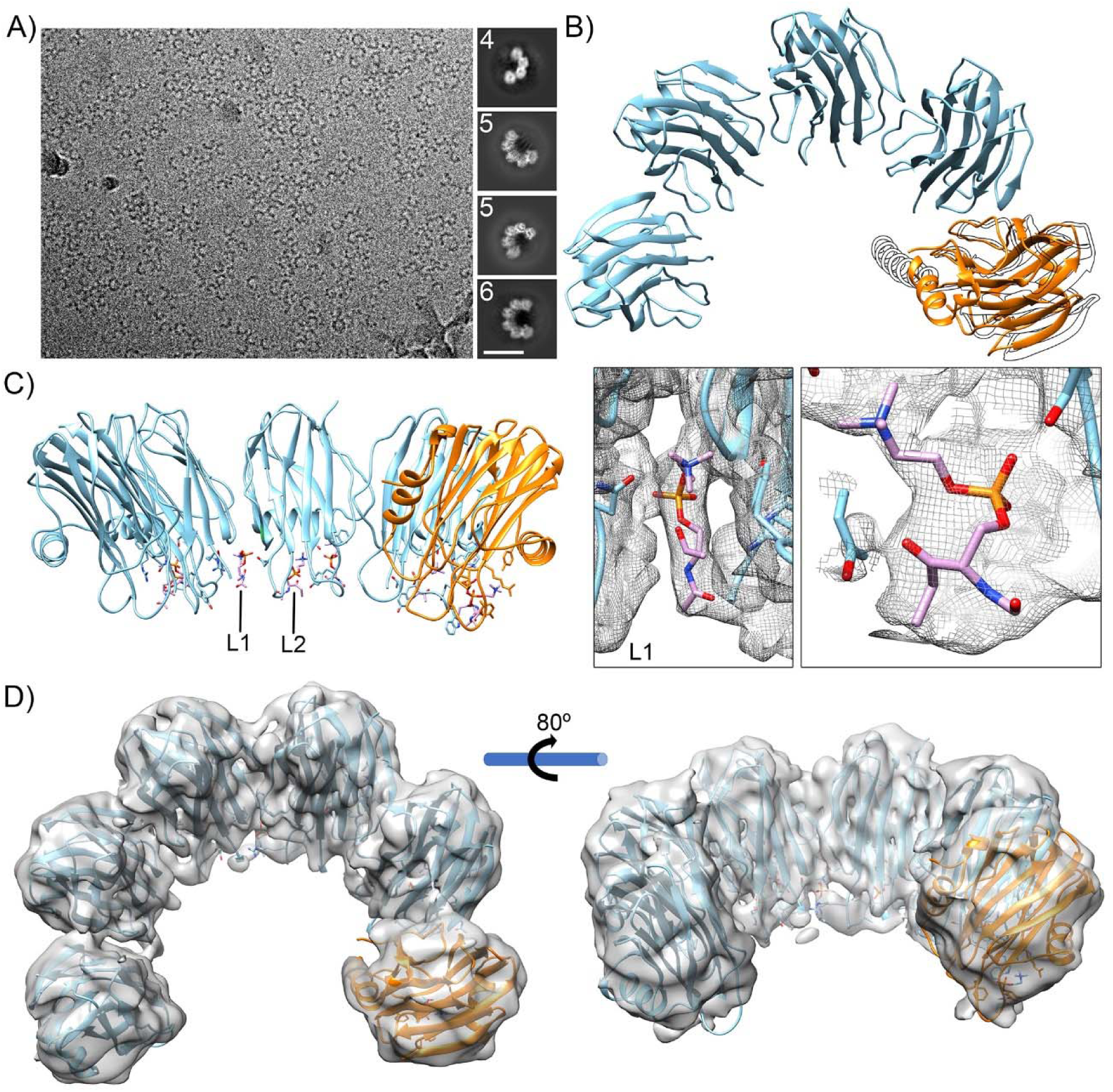
Intermediate pore structures. A) CryoEM image showing lipid nanodiscs containing intermediate pore arrangements (left) alongside 2D averages of structures containing 4, 5, and 6 monomers (right). Scale bar 100 Å. B) Comparison showing the position of the monomer at the end of the arc (brown) with its final position in the complete pore (outlined in black). C) Side view of the intermediate pore showing the positions of the polar head of the lipids (left) and some details of densities corresponding to the SM in the fenestration (L1, center) and lipid 2 (right). This data indicates that some lipids are already associated with the pore even in the early stages of their formation. D) Two views of the map of the 6mer intermediate pore and the atomic structure build on it. As in the 5mer structure the last monomer shows the α helix-1 folded as in soluble form of the protein.

**Extended Data Figure 12.**
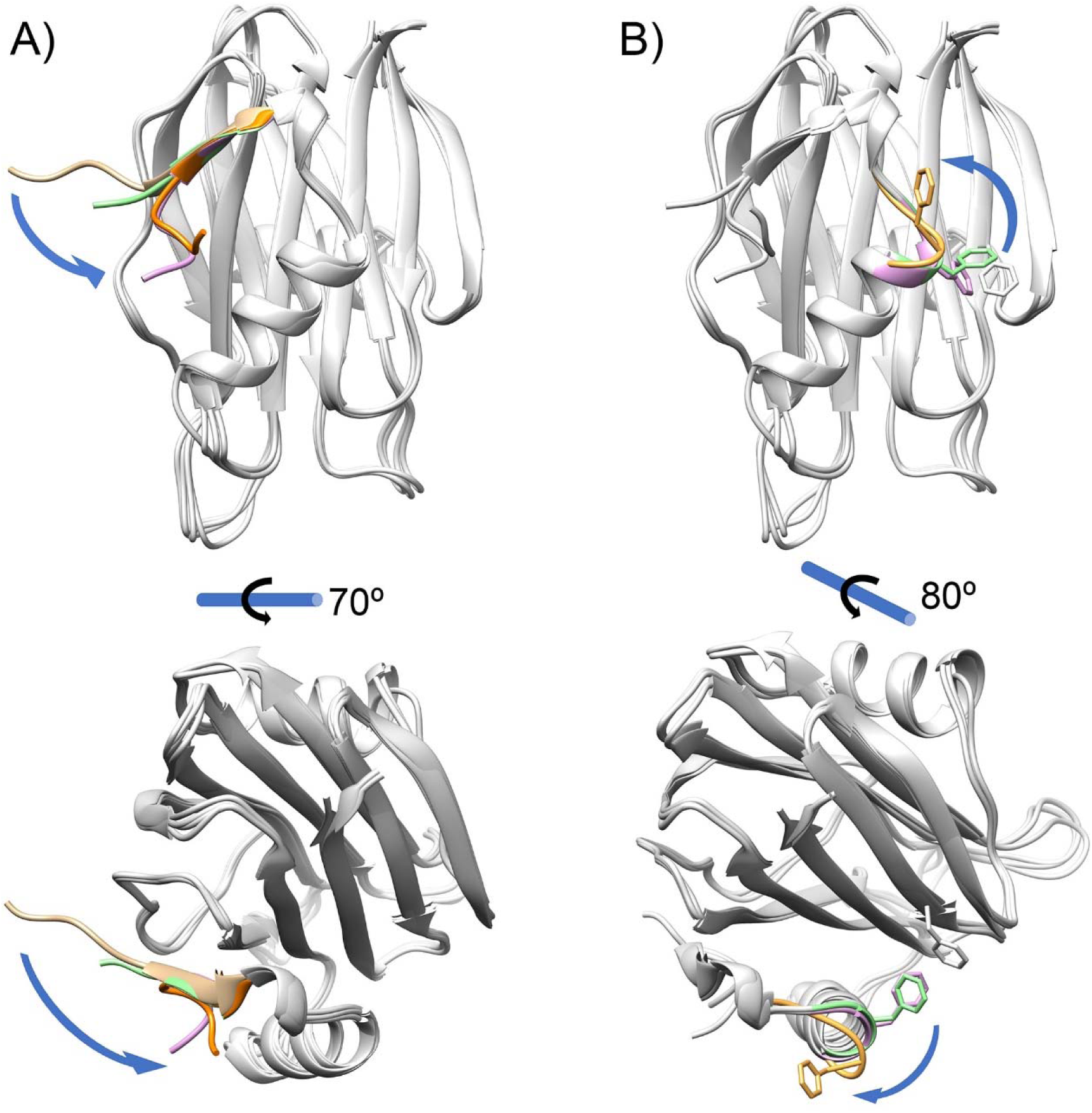
N-terminal/Phe 14 conformations in different atomic PFTs structures. A) N-terminal could adopt different positions. The superposition of crystallographic structures of StnII (pink), FraC in two conformations (light and dark brown) and EqtII (light green) reveals the position variability of the N-terminal. B) The conserved Phe residue in α-helix 1 could be displaced from its hydrophobic environment (Phe 14 in StnII and Phe 16 in FraC and EqtII). StnII is shown in pink, FraC in light brown and EqtII in light green. In both panels, the rest of the molecule has been colored gray for clarity and conformational changes have been depicted with blue arrows. PDB data: For panel A) StnII 1GWY, pink [11]; FraC 4TSQ (chain E) light brown and 3VWI dark brown [13] and EqtII 1TQZ, light green [10]. For panel B) StnII 1GWY, pink [11]; FraC 4TSN light brown [13] and EqtII 1TQZ, light green [10].

## Notes

### Competing Interest Statement

The authors have declared no competing interest.

### Summary of Updates

- Accesion codes for Electron Microscopy Data Bank (EMDB) and Protein Data Bank (PDB). - References.

